# Prefoldin function links meiotic chromosome segregation with cellular remodeling and reveals tubulin sensitivity of the meiotic spindle

**DOI:** 10.64898/2026.05.20.726416

**Authors:** Naohiro Kuwayama, Benjamin S Styler, Bojana Stekovic, Helen Sakharova, Liana Lareau, Marko Jovanovic, Elçin Ünal, Gloria A Brar

## Abstract

Faithful chromosome segregation is essential for producing viable gametes during meiosis, a specialized type of cell division compared to mitosis. Unlike mitosis, meiosis involves two consecutive chromosome segregation events without an intervening round of DNA replication. Here we identify Gim3, a subunit of the ubiquitously expressed and conserved prefoldin complex, as a critical regulator of meiotic but not mitotic chromosome segregation in budding yeast. Loss of Gim3 causes profound defects in chromosome segregation and gamete viability through reduced tubulin protein levels, which are also associated with reduced spindle length. In mitosis, however, *GIM3* deletion minimally affects spindle length and chromosome segregation, despite similarly reduced tubulin levels in both contexts, highlighting a previously unrecognized difference between the sensitivity of meiotic and mitotic spindles to tubulin abundance. In addition to chromosome segregation defects, *gim3Δ* cells exhibit aberrant meiotic cellular remodeling, including defects in exclusion of age-associated protein aggregates from newly forming gametes. Importantly, experimentally induced meiotic chromosome mis-segregation similarly disturbs cellular remodeling. Together, our findings identify Gim3 as a key factor required for maintaining chromosome segregation integrity during meiosis and reveal a previously unrecognized link between chromosome segregation and meiotic cellular remodeling.

## Introduction

In eukaryotes, sexual reproduction is based on the generation of haploid gametes through gametogenesis, a highly regulated developmental program in which diploid progenitor cells undergo meiosis to halve their genome. This cycle is characterized by two rounds of chromosome segregation after a single DNA replication phase followed by homologous recombination. During the first segregation phase, meiosis I (MI), homologous chromosomes are separated. During meiosis II (MII), sister chromatids separate. Pioneering studies in yeast have uncovered conserved mechanisms that ensure faithful chromosome segregation during this process (Nasmyth, 2002; Marston et al., 2004; Watanabe, 2012). These studies have demonstrated that factors such as Rec8, Sgo1, and the monopolin complex, which directly regulate chromosome cohesion and kinetochore orientation, are critical for accurate chromosome segregation during meiosis (Klein et al., 1999; Kitajima et al., 2004; Corbett et al., 2010). Such factors help to overcome the physical challenges of separating recombined homologous chromosomes at MI and removing cohesion between sister chromatids in two stages to ensure the fidelity of chromosome segregation at both MI and MII, features that are not required in mitosis. Beyond these differences, the mechanics of meiotic segregation, especially MII, have been thought to be similar to the segregation that occurs in mitosis (Petronczki et al., 2003; Winey et al., 2005).

In addition to faithful chromosome segregation, production of healthy gametes relies on quality control mechanisms that ensure proper distribution of nuclear and cytoplasmic contents, which occurs by largely distinct mechanisms in meiosis and mitosis (King and Ünal, 2020; Xiao and Ünal, 2025). Mitotic divisions in yeast are asymmetric, and components are either segregated to the daughter cell or retained in the mother cell (Lin et al., 2025). Meiotic divisions, in contrast, are both symmetric and involve destruction of much of the preexisting cellular contents through multiple modes of protein degradation. For example, components of the nuclear pore complex (NPC) are selectively segregated from the gamete nuclei and subsequently degraded (King et al., 2019). Similarly, organelles such as the endoplasmic reticulum (ER) and mitochondria undergo extensive reorganization, potentially to ensure that only functional components are inherited by the gametes (Suda et al., 2007; Sawyer et al., 2019; Otto et al., 2021). These quality control events are thought to be critical in the process of rejuvenation, which resets the lifespan of the next generation, particularly in aged cells (Ünal et al., 2011). Protein aggregates, including those associated with the chaperone Hsp104, which accumulate in aged cells, are segregated and eliminated during gametogenesis (King et al., 2019). Despite the importance of these processes for gamete production and potentially rejuvenation, it is not yet understood how they are regulated at a molecular level and coordinated with major meiotic events like chromosome segregation.

Prefoldin is a conserved chaperone present in organisms ranging from archaea to humans. It was originally identified in budding yeast as the hexameric GimC complex, composed of Gim1-Gim6 in yeast and PFDN1-PFDN6 in humans, and is required for diverse cellular processes (Geissler, 1998; Liang et al., 2020; Tahmaz et al., 2022). Prefoldin transfers clients to the chaperonin complex for further folding, with its best characterized clients being actin and tubulin subunits (Vainberg et al., 1998; Siegers, 1999; Rommelaere et al., 2001). Genetic and functional analyses have demonstrated that it contributes broadly to cellular homeostasis, embryonic development, and lifespan (Delgehyr et al., 2012; Lundin et al., 2008; Son et al., 2018; Zhang et al., 2016). In humans, prefoldin dysfunction has been associated with cancer and neurodegenerative diseases (Herranz-Montoya et al., 2021; Tashiro et al., 2013). Although all the above roles of prefoldin are associated with the conserved complex, there have been reports that individual prefoldin subunits may be capable of acting out of the context of the defined heterohexameric complex to perform separate functions, including regulating gene expression (Rodríguez-Milla and Salinas, 2009; Millán-Zambrano et al., 2013; Payán-Bravo et al., 2021; Rodrigues et al., 2023; Tahmaz et al., 2023; Shahmoradi Ghahe et al., 2024).

Here, through comprehensive screening, and genetic and microscopic analyses, we identified the surprising requirement of a prefoldin subunit, Gim3, in regulating chromosome segregation specifically during meiosis. In this context, Gim3 functions as part of the prefoldin complex, with tubulin as a key client. Our findings reveal that Gim3 is required not only for accurate meiotic chromosome segregation but also for the cellular remodeling that underlies gamete production, including exclusion of age-associated protein aggregates from gametes. Notably, induction of chromosome mis-segregation disrupts cellular remodeling and quality control during meiosis, revealing a previously unrecognized role for chromosome segregation in driving broader cellular remodeling and quality control during meiosis.

## Results

### Gim3 deficiency results in chromosome mis-segregation during meiosis

While investigating the function of non-canonical open reading frames (ORFs) in meiosis using a CRISPR-based screen, we unexpectedly identified Gim3, encoded by a canonical ORF, as a key regulator of meiosis. In brief, we constructed guide RNA libraries targeting most short ORFs (sORFs) and truncated ORFs (tORFs) detected as translated during meiosis (Brar et al., 2012; Higdon et al., 2024), introduced mutations disrupting translation of these non-canonical coding regions into yeast populations, and subsequently induced meiosis and gamete formation (sporulation). Gametes were subsequently enriched by zymolyase treatment to remove non-sporulated cells, and we sequenced the guide RNAs from the non-treated versus treated populations (Figure S1A, File S1). As expected, positive controls targeting coding regions previously reported to be important for meiosis were detected among the highest-scoring hits (Figure S1B). We next examined top-scoring sORF mutants by generating full ORF deletion mutants and tested their efficiency of progression through meiosis. Disruption of “*sORF2*” (positioned on chr14 plus strand, upstream of *YNL152W*) increased the proportion of mono-nucleated cells relative to spores, suggesting that this locus is important for timely progression of meiosis (Figure S1C). Notably, *sORF2* is embedded in the *GIM3* locus, and translated from an antisense transcript to *GIM3* (Figure S1D). To determine whether Gim3 or sORF2 was responsible for the observed phenotype, we conducted rescue experiments. Overexpression of *GIM3*, but not *sORF2*, restored normal sporulation, indicating that Gim3 is critical for this function (Figure S1E). Consistently, deletion of *GIM3* (which also removes *sORF2*) resulted in slower sporulation compared to wild-type (WT) cells (Figure 1A). Importantly, *gim3Δ* cells showed comparable mitotic growth to WT cells (Figure 1B), implying a critical role for Gim3 that is specific to meiosis.

**Figure 1.**
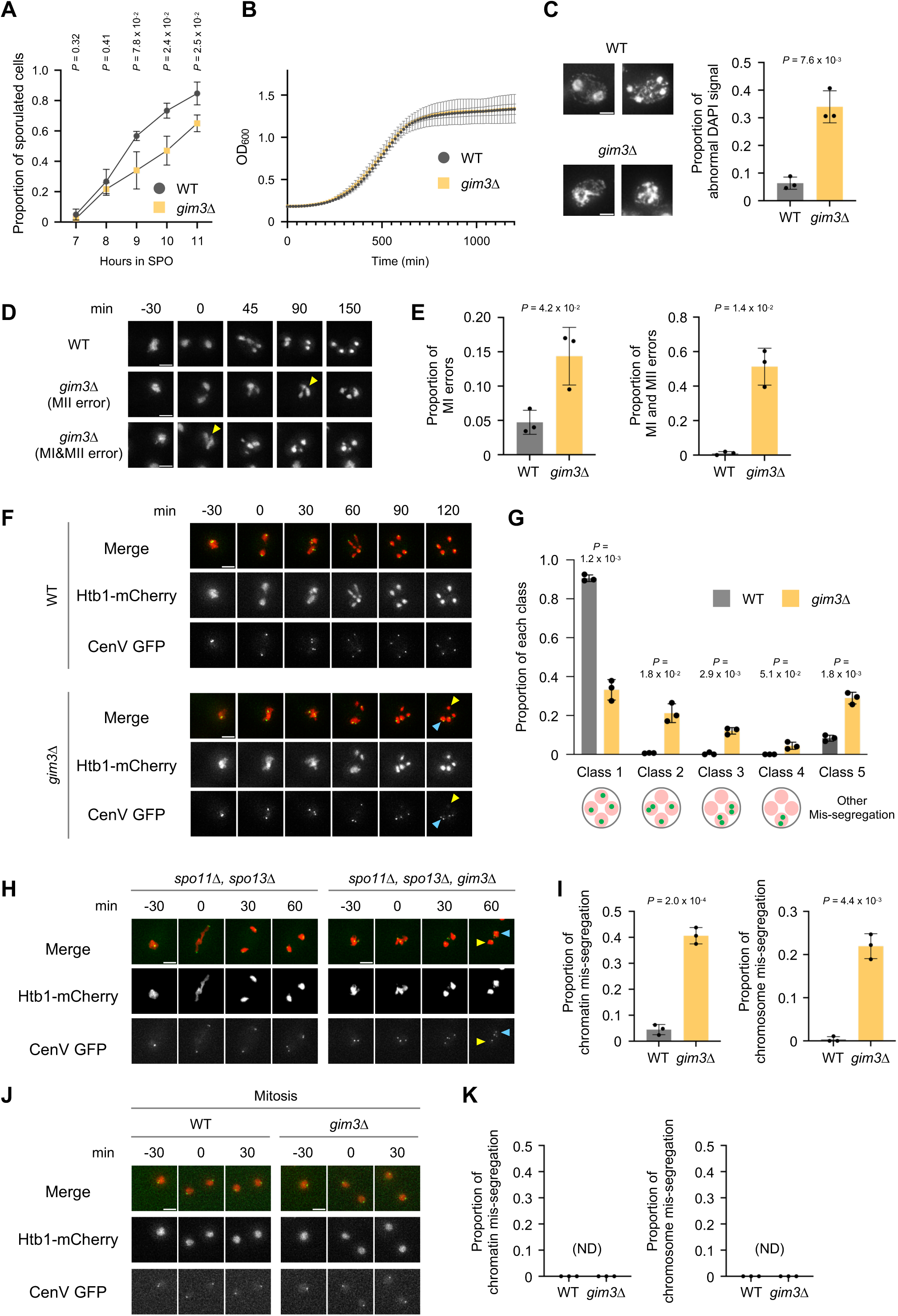
Gim3 deficiency results in chromosome mis-segregation during meiosis. (A) Meiotic progression as determined by DAPI staining for WT and *gim3Δ* strains. N = 3; data are represented as mean ± SD; Welch’s t-test. 100 cells were quantified per replicate. (B) Growth curves of WT and *gim3Δ* strains in YPD, monitored by OD_600_ measurements. N = 3. Data are represented as mean ± SD. (C) Representative images of WT and *gim3Δ* cells at 6 hours in sporulation medium, stained with DAPI. Quantification of abnormal nuclear morphology by DAPI staining is shown. N = 3; data are represented as mean ± SD; Welch’s t-test. 147-209 cells were quantified per replicate. Scale bar, 2 µm. (D) Time-lapse microscopy of cells expressing Htb1-mCherry to mark chromatin imaged every 15 min during meiosis. A WT cell (top), a *gim3Δ* cell showing a MI error (middle), and a *gim3Δ* cell showing MI and MII errors (bottom) are presented. Yellow arrowheads indicate chromatin mis-segregation. Time 0 was set at the onset of anaphase I, when two chromosome masses first pull apart. Scale bars, 3 µm. (E) Quantification of Htb1-mCherry-marked mis-segregation events during MI (left) and MI&MII (right). N = 3; data are represented as mean ± SD; Welch’s t-test. 63-157 cells were quantified per replicate. (F) Live-cell imaging of Htb1-mCherry and CenV GFP in WT (top) and *gim3Δ* (bottom) cells imaged every 15 min during meiosis. Yellow arrowheads indicate chromatin masses lacking a CenV GFP dot, and blue arrowheads indicate a chromatin mass containing two CenV GFP dots. Time 0 was set at the onset of anaphase I. Scale bars, 3 µm. (G) Quantification of chromosome mis-segregation shown in (F). Cells were classified into five classes based on the number of chromatin masses lacking a CenV GFP dot: Class 1, all chromatin masses contain one CenV GFP dot; Class 2, one chromatin mass lacks a CenV GFP dot; Class 3, two chromatin masses lack CenV GFP dots; Class 4, three chromatin masses lack CenV GFP dots; Class 5, other types of chromatin mis-segregation. N = 3 biological replicates; data are presented as mean ± SD; Welch’s t-test. 98-178 cells were quantified per replicate. (H) Live-cell imaging of Htb1-mCherry and CenV GFP dots in *spo11Δ spo13Δ* (control, left) and *spo11Δ spo13Δ gim3Δ* (right) cells. Yellow arrowheads indicate chromatin masses containing three CenV GFP dots, and blue arrowheads indicate a chromatin mass containing one CenV GFP dot. Time 0 was set at the onset of anaphase. Scale bars, 3 µm. (I) Quantification of chromatin (left) and chromosome (right) mis-segregation events from (H). N = 3; data are represented as mean ± SD; Welch’s t-test. 54-126 cells were quantified per replicate. (J) Htb1-mCherry and CenV GFP dot imaging during mitosis in WT and *gim3Δ* cells. Images were taken every 10 min during mitosis. Diploid strains were used for this experiment. Time 0 was set at the onset of anaphase. Scale bars, 3 µm. (K) Quantification of chromatin (left) and chromosome (right) mis-segregation during mitosis from (J). N = 3; data are represented as mean ± SD. ND, not detected. 69-91 cells were quantified per replicate.

Microscopic analysis of DNA stained with DAPI in *gim3Δ* cells during meiosis revealed frequent abnormal patterns, characterized by fuzzy or interconnected DAPI signals that were rare in WT cells at matched stages (Figure 1C). This suggested possible defects in chromosome segregation in *gim3Δ* cells. To further examine this phenotype, we first performed live imaging using a chromatin marker (Htb1-mCherry). We found that *gim3Δ* cells displayed a markedly increased frequency of abnormal chromatin distribution during both MI and MII, typically manifested as incomplete separation of chromatin during MI or uneven nuclear sizes during MII (Figure 1D, E). To further monitor chromosome segregation, we used the TetO/TetR-GFP strategy to homozygously label chromosome V at a site marked with tandem Tet operator sequences inserted 1.4 kb from the centromere (CenV GFP) (Straight et al., 1996; Lee and Amon, 2003). Consistent with the chromatin analysis, *gim3Δ* cells exhibited a significant increase in chromosome mis-segregation of this marked chromosome compared to WT cells, as judged by the presence of more than one CenV GFP dot within a single Htb1-mCherry marked nuclear mass following both MI and MII (Figure 1F). We classified the CenV GFP dot patterns into five classes based on the number of chromatin masses lacking CenV GFP dots. This analysis showed that *gim3Δ* cells exhibited a higher proportion of chromatin masses without CenV GFP dots in multiple patterns (Figure 1G), indicative of mis-segregation during both MI and MII. To determine whether MII mis-segregation in *gim3Δ* cells was secondary to preceding MI errors or reflected an inherent defect in this second stage of chromosome segregation, in which centromeric cohesion is retained and kinetochores are bioriented, we examined *gim3Δ* in a *spo11Δ spo13Δ* background, in which recombination is inhibited, and cells undergo a single MII-like division (Lee and Amon, 2003). In this background, *gim3Δ* cells showed elevated levels of mis-segregation for both Htb1-mCherry and CenV GFP, indicating that Gim3 is intrinsically required for accurate chromosome segregation at MII as well as MI (Figure 1H, I). Importantly, no defects in chromosome segregation were observed during mitosis (Figure 1J, K), indicating that Gim3 function in supporting chromosome segregation is specifically critical for meiosis.

Sporulation medium (SPO) differs in composition substantially from the rich media (YPD) that is typically used for mitotic growth. It contains acetate rather than a fermentable carbon source, inducing respiration, and also lacks yeast extract, which includes an array of poorly defined growth-enhancing components. We tested mitotic growth in pre-sporulation medium (BYTA), which contains acetate, as well as synthetic medium, which lacks yeast extract. Chromosome mis-segregation was not detected in either condition (Figure S2A, B). As a complementary approach, we took advantage of an approach showing that loss of protein kinase A (PKA) activity through pharmacological inhibition of the yeast PKA *TPK1* (through use of the *tpk1-as* mutant) in cells lacking the other two PKA-encoding genes, *TPK2* and *TPK3*, together with inhibition of TORC1 by rapamycin, allows induction of meiosis even in YPD (Weidberg et al., 2016). We asked whether *gim3Δ* cells exhibit chromosome segregation defects in YPD under these conditions. Indeed, *gim3Δ tpk1-as tpk2Δ tpk3Δ* cells treated with 1NM-PP1 and rapamycin displayed an increased frequency of chromosome mis-segregation relative to *tpk1-as tpk2Δ tpk3Δ* cells with Gim3, with matched treatment conditions, after MII in this rich media context (Figure S2C, D). These results indicate that the chromosome segregation defect in *gim3Δ* cells is inherent to the meiotic chromosome segregation program and not simply caused by media dependent changes in cellular state. Together, these findings establish Gim3 as a factor required for faithful chromosome segregation during meiosis, a novel role for this conserved factor.

### Loss of Gim3 or other prefoldin components impairs spore viability

Faithful meiotic chromosome segregation is essential for producing healthy gametes. To assess whether *gim3Δ* affects the viability of gametes, we dissected tetrads from *gim3Δ* diploids and observed a significant reduction in spore viability compared with WT controls (Figure 2A, B). This finding is consistent with the rampant chromosome mis-segregation observed in these cells (Figure 1C-I). Importantly, reintroduction of *GIM3*, but not *sORF2*, prior to meiosis rescued this spore viability defect (Figure 2C, D, S1F), underscoring the critical role of Gim3 in ensuring the production of viable gametes. One possible explanation for the low viability of *gim3Δ* spores could be that they are impaired in germination, or growth, following meiosis. To address this, we dissected tetrads produced from diploid cells that were heterozygous for *gim3Δ*. If *gim3Δ* impaired post-dissection growth, spores that inherit this allele would show reduced growth compared with their WT counterparts. However, growth of *gim3Δ* haploids was indistinguishable from control in this condition (Figure 2E, F), indicating that the reduced viability originates from Gim3 function during meiosis.

**Figure 2.**
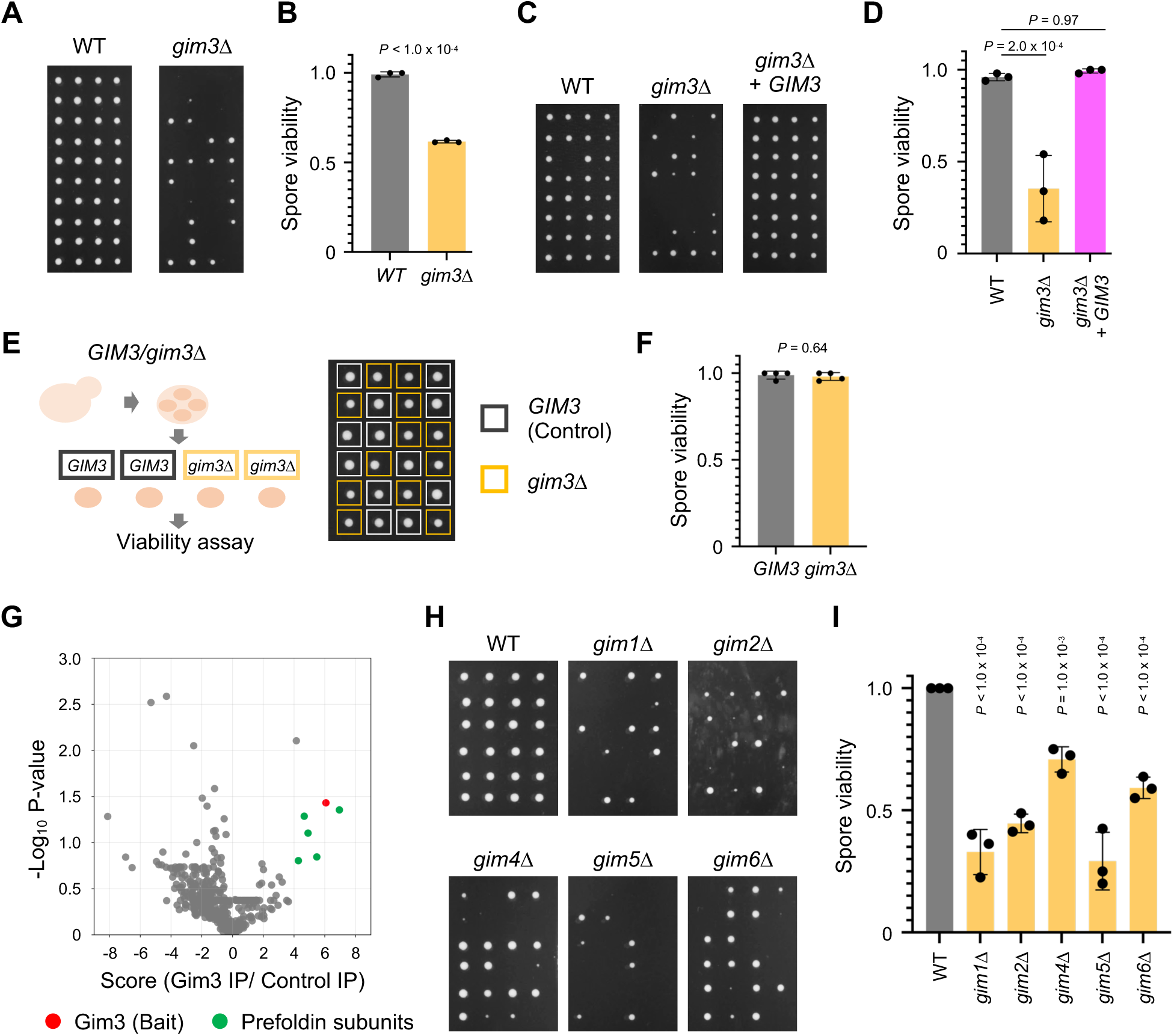
Loss of Gim3 or other prefoldin components impairs spore viability. (A) Spore colony growth of tetrads derived from WT and *gim3Δ* strains on YPD after 2 days at 30 °C. (B) Quantification of spore viability from (A). N = 3; data are represented as mean ± SD; Welch’ s t-test. 80 spores were quantified per replicate. (C) Spore colony growth of WT, *gim3Δ* with empty cassette, and *gim3Δ* with *GIM3* complementation after 2 days at 30 °C. (D) Quantification of spore viability from (C). N = 3; data are represented as mean ± SD. Dunnett’s multiple comparison test. 64-80 spores were quantified per replicate. (E) Schematic of heterozygous diploids (*GIM3*/*gim3Δ*) used in a spore viability assay (left). Spore colony growth of tetrads derived from *GIM3/gim3Δ* strains on YPD after 2 days at 30 °C (right). Genotypes were determined after replica plating on G418 plates. (F) Quantification of spore viability from (E). N = 3; Welch’s t-test; data are represented as mean ± SD. 44-64 spores were quantified per replicate. (G) Immunopurification-mass spectrometry (IP-MS) analysis of Gim3-3V5 cells collected after 5 hours in SPO. Plot shows enrichment (x-axis) versus –log_10_ P-value (y-axis). Gim3 (bait) is indicated in red; prefoldin subunits are indicated in green. (H) Spore colony growth of tetrads derived from WT and individual prefoldin subunit deletion strains (*gim1Δ, gim2Δ, gim4Δ, gim5Δ, gim6Δ*). (I) Quantification of spore viability from (H). N = 3; data are represented as mean ± SD. Dunnett’s multiple comparison test. 76-80 spores were quantified per replicate.

We next asked how Gim3 contributes to chromosome segregation and spore viability. Previous studies suggested that Gim3 can act as a co-chaperone within the prefoldin complex, but also that individual subunits may interact with other factors to regulate transcription or proteasome assembly (reviewed in Liang et al., 2020; Tahmaz et al., 2022). To distinguish these potential modes of action, we performed immunopurification-mass spectrometry (IP-MS) using Gim3-3V5 as bait during meiosis as well as mitosis. Factors enriched in the Gim3 IP included all other prefoldin subunits (Figure 2G, S3A-C, File S2). Furthermore, Cct4, a subunit of the chaperonin CCT that receives clients from prefoldin, was also present in Gim3 IP samples (Figure S3C). These results suggest that Gim3 functions within the prefoldin complex during meiosis. If the critical function of Gim3 in meiosis is based on its role as part of the prefoldin complex, then deletion of other prefoldin subunits should also impair spore viability. Consistent with this idea, deletion of the genes encoding Gim1, Gim2, Gim4, Gim5, or Gim6 reduced spore viability without grossly affecting mitotic growth (Figure 2H, I, S3D). Together, these results therefore identify a function of the entire prefoldin complex that is important to support normal production of spores.

### Gim3 is required for maintaining tubulin levels and proper spindle formation

Given that prefoldin serves as a co-chaperone, delivering client proteins to chaperonin, we next sought to identify important meiotic client proteins of Gim3. To this end, we performed mass spectrometry on meiotic lysates from *gim3Δ* cells and compared protein abundances with WT cells. Gene ontology analysis revealed that proteins reduced in *gim3Δ* were significantly enriched for functional categories related to tubulin and actin (Figure 3A, B, File S3), supporting a model in which these well-characterized prefoldin clients could be the major clients in meiosis. Indeed, Tub1 protein levels were consistently lower in *gim3Δ* cells during the meiotic time course (3-6 hours in SPO) than in WT cells (Figure 3C). Western blot analysis further validated the reduction in Tub1 in both mitosis and meiosis *gim3Δ* cells (Figure 3D, E, S4D). Ribosome profiling indicated that *TUB1* translation levels were similar between WT and *gim3Δ* cells at 4 hours in SPO (Figure S4A, File S4), suggesting that the reduction in Tub1 levels occurs post-translationally. Moreover, deletion of other prefoldin subunits led to decreased Tub1 levels in both mitosis and meiosis (Figure S4B, C), supporting the notion that Tub1 is a client of the prefoldin complex in both contexts.

**Figure 3.**
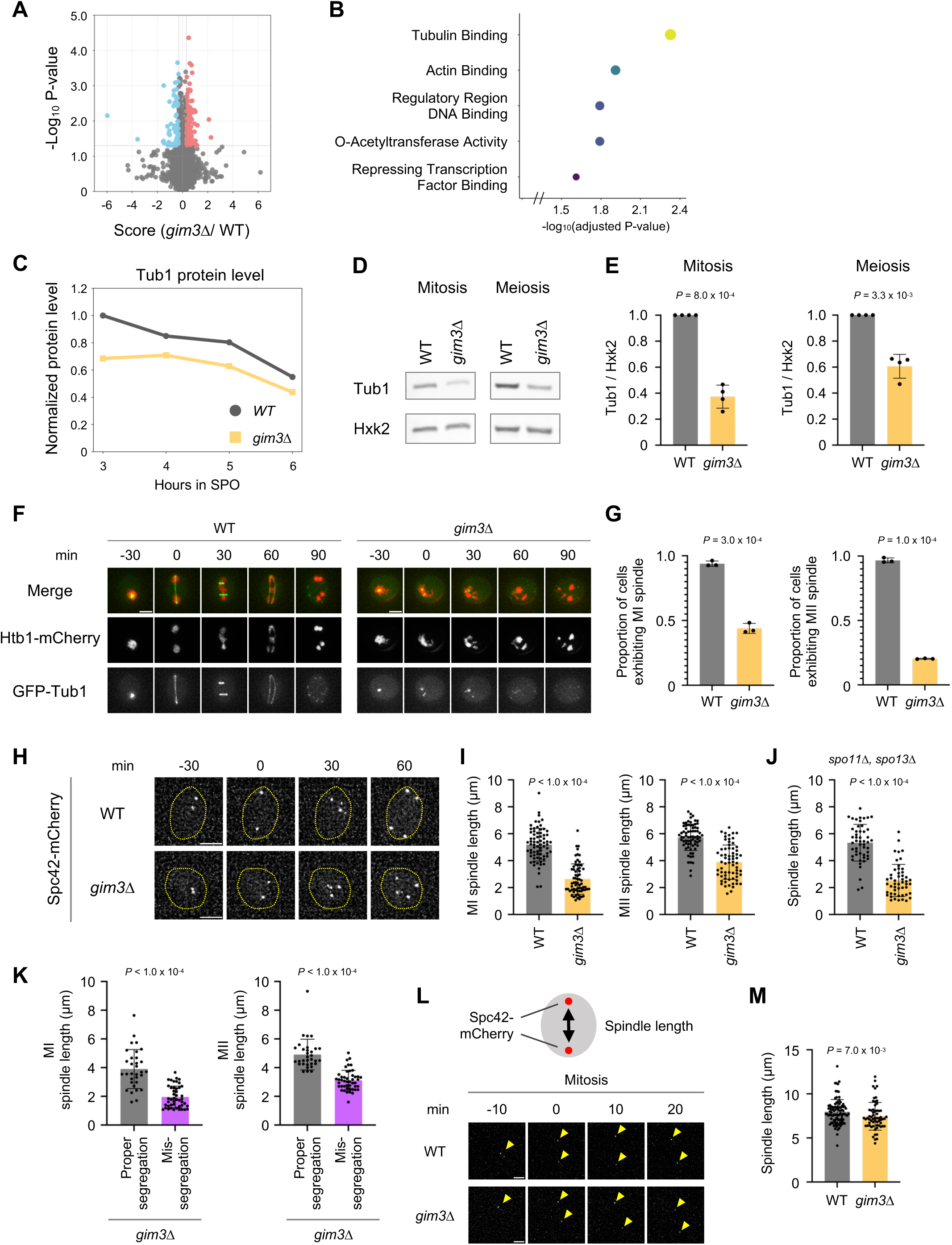
Gim3 is required for maintaining tubulin levels and proper spindle formation. (A) Mass spectrometry analysis comparing WT and *gim3Δ* cells collected after 3-6 hours in SPO. Plot shows enrichment (x-axis) versus –log_10_ P-value (y-axis). Proteins downregulated in *gim3Δ* cells are highlighted in blue, and proteins upregulated in *gim3Δ* cells are highlighted in red. (B) Gene ontology enrichment analysis of proteins downregulated in *gim3Δ* cells, with adjusted P-values indicated. (C) Normalized Tub1 protein levels in WT and *gim3Δ* cells at 3, 4, 5 and 6 hours in SPO, quantified by mass spectrometry. Mass spectrometry analysis was performed once for each time point. (D) Representative western blot of Tub1 in WT and *gim3Δ* cells during exponential mitotic growth and after 4 hours in SPO (meiosis). Hxk2 (Hexokinase isoenzyme 2) was used as a loading control. (E) Quantification of Tub1 protein levels from (D). N = 4; data are represented as mean ± SD; Welch’s t-test. (F) Live-cell imaging of WT and *gim3Δ* cells expressing Htb1-mCherry and GFP-Tub1 during meiosis. Time is shown relative to the onset of anaphase I (0 min). Anaphase II spindles can be seen at 60 min. Scale bars, 3 µm. (G) Quantification of the proportion of cells exhibiting anaphase I (left) and anaphase II (right) spindles among cells followed through meiotic divisions in the live-cell imaging experiment shown in (F). N = 3; data are represented as mean ± SD; Welch’s t-test. 54-120 cells were quantified per replicate. (H) Live-cell imaging of WT and *gim3Δ* cells expressing Spc42-mCherry to mark spindle pole bodies during meiosis. Time is shown relative to the onset of anaphase I (0 min). Scale bars, 3 µm. (I) Quantification of anaphase I and anaphase II spindle length from (H). Each dot represents an individual cell. Bars indicate the mean ± SD. Mann-Whitney U test; 67-72 cells were quantified. (J) Spindle length at anaphase I in *spo11Δ spo13Δ* and *spo11Δ spo13Δ gim3Δ* cells expressing Spc42-mCherry. Each dot represents an individual cell. Bars indicate the mean ± SD. Mann-Whitney U test; 50 cells were quantified. (K) Spindle lengths at anaphase I (left) and anaphase II (right) in *gim3Δ* cells showing either proper chromatin segregation or mis-segregation. N = 3; data are presented as mean ± SD; Welch’s t-test. 25-27 cells were quantified per replicate. (L) Schematic and live imaging of Spc42-mCherry during mitosis in WT and *gim3*Δ cells. Images were taken every 10 min during mitosis. Time 0 was set at the onset of anaphase. Scale bar, 2 µm. Yellow arrowheads indicate Spc42-mCherry signals. (M) Quantification of spindle length at anaphase from (L). N = 3; data are represented as mean ± SD; Welch’s t-test. 37-53 cells were quantified per replicate.

Because tubulin is the primary component of the spindle microtubules, we analyzed spindle morphology using GFP-tagged Tub1 in cells with or without Gim3. In WT cells, elongated anaphase spindles were readily observed during both MI and MII, whereas *gim3Δ* cells exhibited a markedly lower proportion of cells with morphologically normal spindles (Figure 3F, G). To more directly estimate microtubule polymer mass in the meiotic spindle, we quantified the fluorescence intensity of GFP-Tub1 within the spindle. GFP-Tub1 signal intensity was reduced in *gim3Δ* cells compared with WT, suggesting decreased spindle microtubule polymer mass (Figure S4E). In addition, we noted that the length of anaphase spindles appeared shorter in cells lacking Gim3 than WT cells. We assessed spindle length using measurement of the maximum distance between foci for the spindle pole body marker Spc42 (Miller et al., 2012). Spindle length was significantly reduced in *gim3Δ* cells during both MI and MII (Figure 3H, I). Notably, this phenotype was also observed in the *spo11Δ spo13Δ* background, indicating that the defect is independent of recombination and occurs for both MI and MII-like spindles (Figure 3J). Notably, among *gim3Δ* cells, those exhibiting chromosome mis-segregation had much shorter spindle lengths compared to cells without mis-segregation, suggesting that chromosome segregation errors are associated with reduced spindle length (Figure 3K). By contrast, spindle length during mitosis showed only a modest reduction in *gim3Δ* cells compared to WT (8.0% ± 4.0%; Figure 3L, M), which was markedly smaller than the reductions observed during meiosis I (48.9% ± 6.1%, Figure 3H, I) and meiosis II (37.6% ± 3.9%, Figure 3H, I), despite similar tubulin abundance in mitosis and meiosis (Figure 3D). Together, our results demonstrate that prefoldin maintains proper tubulin protein levels in both mitotic and meiotic cells, but that loss of Gim3 impairs spindle elongation preferentially during meiosis, potentially through higher sensitivity of the meiotic spindles to tubulin insufficiency.

Chromosome segregation depends on both proper spindle elongation and attachment of chromosomes to spindle microtubules through kinetochores. Aberrant chromosome-spindle attachments can prevent spindle elongation. Defective attachment between chromosomes and spindle microtubules activates the spindle assembly checkpoint (SAC), leading to a delay in cohesin cleavage and spindle elongation to allow time for correct attachments to occur (Musacchio and Salmon, 2007). To assess whether meiotic SAC activation in cells lacking Gim3 contributes to spindle elongation defects, we examined the effect of deleting *MAD2*, the gene encoding a key conserved SAC component (Li and Murray, 1991), in *gim3Δ* cells. If chromosome-spindle attachment was strongly impaired by loss of Gim3 function, loss of Mad2 would be expected to exacerbate the segregation defects in cells lacking Gim3. However, *gim3Δ mad2Δ* cells exhibited a level of meiotic chromosome mis-segregation comparable to that of *gim3Δ* cells (Figure S5A). Also consistent with the model that spindle checkpoint hyperactivation does not occur in *gim3Δ* cells, the cleavage of Rec8, the meiotic subunit of the cohesion complex, occurs with similar timing in WT and *gim3Δ* cells, indicating that cohesin removal proceeds normally in the absence of Gim3 (Figure S5B, C). Together, these results suggest that the high rate of chromosome mis-segregation observed in *gim3Δ* cells may be due to insufficient spindle elongation in meiosis arising from reduced tubulin levels rather than from defects in microtubule-kinetochore attachment or cohesin cleavage.

### Reduced Tub1 levels or inhibition of tubulin polymerization leads to chromosome mis-segregation

If reduced tubulin levels are responsible for the meiotic defects observed in *gim3Δ* cells, then reducing tubulin alone should phenocopy the effects associated with loss of Gim3 in meiosis. To test whether this was the case, we examined *TUB1/tub1Δ* heterozygous diploid cells. Tub1 levels in this strain were comparable to those in *gim3Δ* cells during mitosis and were modestly lower during meiosis (Figure S6A, B). Importantly, live imaging of Htb1-mCherry in *TUB1/tub1Δ* cells revealed phenotypes closely resembling those of *gim3Δ* cells, including diffuse chromatin masses and abnormally small nuclei, indicating chromosome segregation errors (Figure 4A, B). CenV GFP dot visualization in cells carrying the TetO/TetR-GFP system corroborated these findings, showing instances in which two or more CenV dots were associated with a single Htb1-mCherry signal in *TUB1/tub1Δ* cells following MII (Figure 4A, B). In contrast, *TUB1/tub1Δ* cells did not exhibit detectable chromosome mis-segregation during mitotic growth in YPD, BYTA, or synthetic media (Figure S6C-E), indicating differential requirements for tubulin levels in mitosis and meiosis. Furthermore, measurement of spindle length using the Spc42 marker demonstrated a significant shortening of both MI and MII spindles in *TUB1/tub1Δ* cells, similar to the phenotype of *gim3Δ* cells (Figure 4C). Consistent with these defects, *TUB1/tub1Δ* cells exhibited substantially reduced spore viability (Figure S6F). While reduced viability is expected given the essential nature of *TUB1*, we observed viability that was much less than half of WT, indicating an additional contribution from chromosome mis-segregation. These data are consistent with a model in which Gim3 function in supporting meiotic spindle elongation is based on its role in supporting normal tubulin abundance.

**Figure 4.**
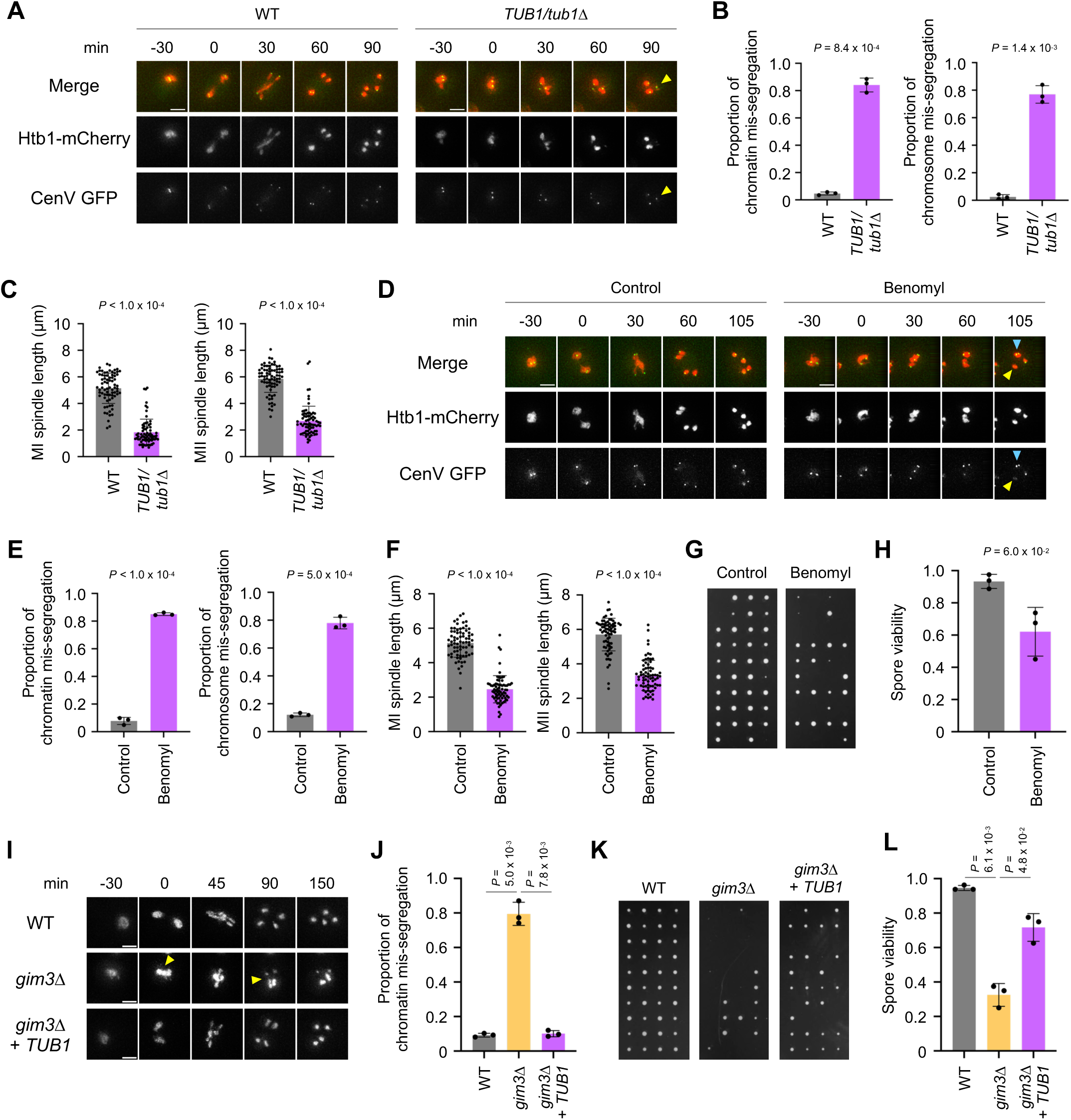
Reduced Tub1 levels or inhibition of tubulin polymerization leads to chromosome mis-segregation. (A) Live-cell imaging of WT and *TUB1/tub1Δ* strains expressing Htb1-mCherry and CenV GFP dots during meiosis. Yellow arrowheads indicate a CenV GFP dot that does not overlap with the corresponding Htb1-mCherry signal. Time 0 was set at the onset of anaphase I. Scale bars, 3 µm. (B) Quantification of mis-segregation events from (A), scored by Htb1-mCherry (left) and CenV GFP (right). N = 3; data are represented as mean ± SD; Welch’s t-test. 56-145 cells were quantified per replicate. (C) Quantification of spindle length at anaphase I (left) and anaphase II (right) in WT and *TUB1/tub1Δ* cells, using Spc42-mCherry to mark spindle pole bodies. Each dot represents an individual cell. Bars indicate the mean ± SD. Mann-Whitney U test; 73-75 cells were quantified. (D) Live-cell imaging of control and benomyl-treated cells expressing Htb1-mCherry and CenV GFP during meiosis. Benomyl was used at 20 μg/mL. Yellow arrowheads indicate chromatin masses lacking a CenV GFP dot, and blue arrowheads indicate a chromatin mass containing two CenV GFP dots. Time 0 was set at the onset of anaphase I. Scale bars, 3 µm. (E) Quantification of mis-segregation events from (D), scored by Htb1-mCherry (left) and CenV GFP (right). N = 3; data are represented as mean ± SD; Welch’s t-test. 58-93 cells were quantified per replicate. (F) Quantification of spindle length at anaphase I (left) and anaphase II (right) in control and benomyl-treated cells, using Spc42-mCherry to mark spindle pole bodies. Each dot represents an individual cell. Bars indicate the mean ± SD. Mann-Whitney U test; 72 cells were quantified. (G) Spore colony growth of tetrads derived from control and benomyl-treated cells. (H) Quantification of spore viability from (G). Data are represented as mean ± SD; N = 3; Welch’s t-test. 80 cells were quantified per replicate. (I) Live-cell imaging of WT cells, *gim3Δ* cells and *gim3Δ* cells expressing *TUB1*, all expressing Htb1-mCherry. Tub1 expression was driven by a GAL promoter and induced using the Gal4-ER system upon addition of β-estradiol (Gao and Pinkham, 2000). β-estradiol was added at 0 hour in SPO to a final concentration of 5 nM to induce expression. Yellow arrowheads indicate chromatin mis-segregation. Time 0 was set at the onset of anaphase I. Scale bars, 3 µm. (J) Quantification of mis-segregation events from (I), scored by Htb1-mCherry. N = 3; data are represented as mean ± SD; Tukey’s multiple comparison test. 129-199 cells were quantified per replicate. (K) Spore colony growth of WT, *gim3Δ* with empty cassette, and *gim3Δ* with *TUB1* complementation after 2 days at 30 °C. (L) Quantification of spore viability from (K). N = 3; data are represented as mean ± SD. Tukey’s multiple comparison test. 80 spores were quantified per replicate.

To determine whether perturbing tubulin dynamics alone is sufficient to cause these defects, we treated meiotic cells with 20 µg/mL of benomyl (Hochwagen et al., 2005), a well-characterized microtubule-destabilizing drug. Strikingly, benomyl-treated cells exhibited the same spectrum of abnormalities as *gim3Δ* and *TUB1/tub1Δ* cells, including diffuse Htb1-mCherry signal, irregular nuclear morphology, multiple CenV GFP signals per Htb1-mCherry signal, and shortened spindles (Figure 4D-F). Benomyl-treated cells also showed reduced spore viability as is observed in *gim3Δ* cells (Figure 4G, H). Lastly, to test whether restoring Tub1 abundance is sufficient to rescue the chromosome mis-segregation phenotype in *gim3Δ* cells, we ectopically induced *TUB1* expression during meiosis (Figure S6G, H). Notably, induction of *TUB1* significantly suppressed meiotic chromosome mis-segregation and restored spore viability in *gim3Δ* cells (Figure 4I-L), demonstrating that restoration of Tub1 abundance rescues the segregation and spore viability defects. In contrast, although Act1 is also a well-characterized prefoldin substrate and was reduced in *gim3Δ* meiotic cells in our conditions (Figure S6I; Rommelaere et al., 2001), ectopic induction of *ACT1* did not rescue the chromosome mis-segregation phenotype or spore viability defects of *gim3Δ* cells (Figure S6J, K). While this does not rule out contributions from other Gim3 substrates, together these experiments support the model that Tub1 is a key Gim3 substrate that explains the meiotic chromosome segregation defects observed in the absence of this prefoldin subunit. The strong rescue from additional Tub1 expression alone was unexpected, as α– and β-tubulin function as an obligate heterodimer, and we therefore anticipated that increasing the abundance of only one subunit would be insufficient to restore spindle integrity. Notably, Tub2 protein levels remained largely unchanged in *gim3Δ* cells, whereas Tub1 levels were substantially reduced, potentially creating an imbalance between α– and β-tubulin abundance (Figure S6L). Because excess orphan β-tubulin has previously been reported to be toxic in yeast (Burke et al., 1989; Weinstein and Solomon, 1990), we considered the possibility that this imbalance contributes to the observed meiotic defects. Chromosome mis-segregation remained substantially elevated in *gim3Δ TUB2/tub2Δ* cells compared to WT controls, indicating that α/β-tubulin imbalance alone cannot account for the meiotic defects caused by loss of Gim3 (Figure S6M). Interestingly, reducing *TUB2* gene dosage in this manner did modestly reduce chromosome mis-segregation in *gim3Δ* cells, suggesting that this imbalance may contribute to the meiotic defects in cells lacking Gim3 (Figure S6M). Together, these findings suggest that insufficient α-tubulin availability is a major cause of the meiotic defects in *gim3Δ* cells. In addition, these experiments revealed a surprising sensitivity to reduced tubulin levels for meiotic, but not mitotic cells.

### Gim3 supports meiotic cellular remodeling and reveals coupling to chromosome segregation

Chromosome segregation during meiosis occurs in a tightly regulated temporal manner that is coordinated with the remodeling of other cellular components. This remodeling ensures the proper inheritance of nuclear and cytoplasmic components while excluding markers of cellular damage, including protein aggregates. One such meiotic cellular remodeling event is the sequestration of pre-existing nuclear pore components, including Nup170, to the Gametogenesis-Uninherited Nuclear Compartment (GUNC), which is not inherited by spores (King et al., 2019). Consistent with previous findings, we saw that Nup170-GFP localized to the GUNC and was subsequently degraded after meiosis in WT cells, reflecting nuclear envelope remodeling (Figure 5A, B). By contrast, in *gim3Δ* cells, Nup170-GFP was less efficiently localized to the GUNC and instead often co-segregated with the chromatin (Figure 5A, B). Consistently, in *gim3Δ* cells, a subset of the Nup170-GFP signal which failed to be sequestered remained in proximity to chromatin, segregating with nuclear masses as meiosis progressed in a pattern that was rarely observed in WT cells (Figure 5A, B). This observation suggests that Gim3 contributes to meiotic cellular remodeling beyond chromosome segregation. These Nup170-GFP-retaining nuclear masses were frequently not packaged into mature spores and instead appeared to be eliminated (Figure S7A-C), reminiscent of the lack of cellularization and aberrant NPC component sequestration observed when the prospore membranes, which ultimately encompass mature gametes, fail to properly form (King et al., 2019). Heterozygous deletion of *TUB1* or treatment of cells with benomyl also resulted in a lower proportion of Nup170 localization to the GUNC, further supporting the idea that proper tubulin dynamics are required for normal meiotic nuclear envelope remodeling (Figure 5C-E, S7D). Consistent with this interpretation, overexpression of *TUB1* in *gim3Δ* cells restored Nup170-GFP sequestration to the GUNC (Figure 5A,B), whereas overexpression of *ACT1* did not, supporting Tub1 as a key substrate linking Gim3 to meiotic nuclear envelope remodeling (Figure S7E). Notably, even with reduced *TUB2* dosage by heterozygous deletion in diploid cells, *gim3Δ* cells failed to efficiently sequester Nup170-GFP (Figure S7F). This result suggests that imbalance between α– and β-tubulin is not sufficient to explain the remodeling defects observed in *gim3Δ* cells. Organelles are also remodeled during meiosis to regulate their inheritance by spores (Suda et al., 2007; Sawyer et al., 2019; Otto et al., 2021). Mitochondria detached from the cell periphery in WT cells, and this also occurred in *gim3Δ* cells (Figure S7G). The cortical ER coalesces into bright, highly dynamic rope-like structures before the first nuclear division and subsequently detaches from the cortex at MII in WT cells. *gim3Δ* cells exhibited normal cortical ER detachment at MII, but showed a markedly reduced frequency of these rope-like structures, the functional significance of which remains unknown (Figure S7H).

**Figure 5.**
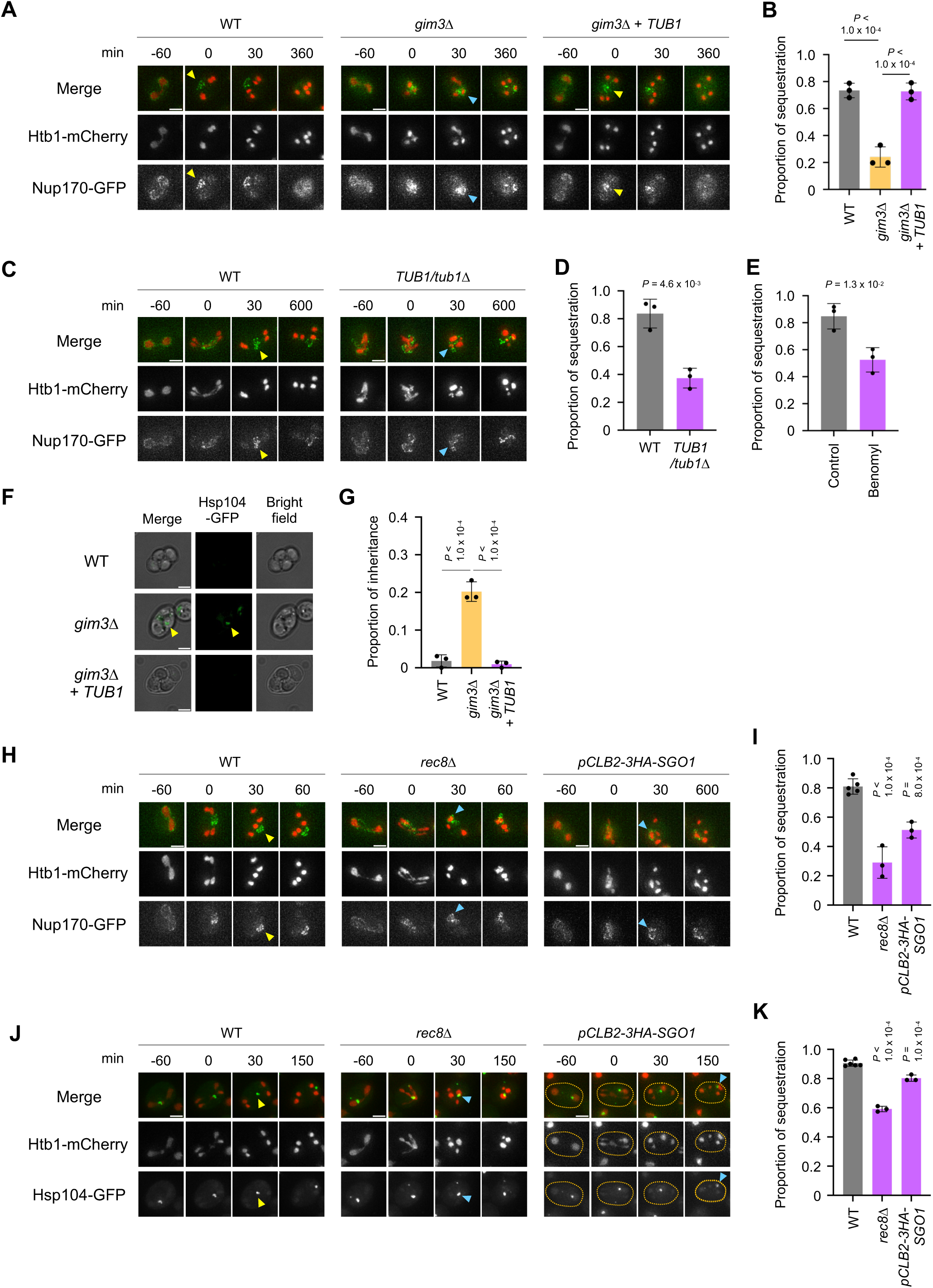
Gim3 supports meiotic cellular remodeling and reveals coupling to chromosome segregation. (A) Live-cell imaging of WT cells, *gim3Δ* cells and *gim3Δ* cells expressing *TUB1*, all expressing Htb1-mCherry and Nup170-GFP during meiosis. Yellow arrowheads indicate sequestration of Nup170-GFP from chromatin, and blue arrowheads indicate co-segregation of Nup170-GFP with chromatin. Time is shown relative to the onset of anaphase II (0 min). Scale bars, 3 µm. (B) Quantification of Nup170-GFP sequestration shown in (A). N = 3; data are represented as mean ± SD; Tukey’s multiple comparison test. 63-88 cells were quantified per replicate. (C) Live-cell imaging of WT and *TUB1/tub1Δ* cells expressing Htb1-mCherry and Nup170-GFP during meiosis. Yellow arrowheads indicate sequestration of Nup170-GFP from chromatin, and blue arrowheads indicate co-segregation of Nup170-GFP with chromatin. Time is shown relative to the onset of anaphase II (0 min). Scale bars, 3 µm. (D) Quantification of Nup170-GFP sequestration shown in (C). N = 3; data are represented as mean ± SD; Welch’s t-test. 49-117 cells were quantified per replicate. (E) Quantification of Nup170-GFP sequestration under benomyl treatment. Benomyl was used at 20 μg/mL. N = 3; data are represented as mean ± SD; Welch’s t-test. 65-123 cells were quantified per replicate. (F) Representative images of WT cells, *gim3Δ* cells and *gim3Δ* cells expressing *TUB1*, all expressing Hsp104-GFP after 22 hours in SPO. Yellow arrowheads indicate Hsp104-GFP signal packaged into spore. Scale bars, 3 µm. Images represent z-slices. (G) Quantification of Hsp104-GFP inheritance from (F). N = 3; data are represented as mean ± SD; Tukey’s multiple comparison test. 64-99 cells were quantified per replicate. (H) Live-cell imaging of WT, *rec8Δ* and *pCLB2-3HA-SGO1* cells, all expressing Htb1-mCherry and Nup170-GFP during meiosis. Yellow arrowheads indicate sequestration of Nup170-GFP, and blue arrowheads indicate co-segregation of Nup170-GFP with chromatin. Time is shown relative to the onset of anaphase II (0 min). Scale bars, 3 µm. (I) Quantification of Nup170-GFP sequestration from chromatin shown in (H). N = 3; data are represented as mean ± SD; Dunnett’s test. 53-126 cells were quantified per replicate. (J) Live-cell imaging of aged WT, *rec8Δ* and *pCLB2-3HA-SGO1* cells, all expressing Htb1-mCherry and Hsp104-GFP during meiosis. Yellow arrowheads indicate sequestration of Hsp104-GFP from chromatin, and blue arrowheads indicate co-segregation of Hsp104-GFP with chromatin. Time is shown relative to the onset of anaphase II (0 min). Scale bars, 3 µm (K) Quantification of Hsp104-GFP sequestration from chromatin shown in (J). N = 3-6; data are represented as mean ± SD; Dunnett’s test. 79-168 cells were quantified per replicate.

Exclusion of protein aggregates from spores, as determined by visualization of Hsp104-GFP foci, occurs in WT cells and has been associated with gametogenic quality control. These protein aggregates are observed in aged precursor cells but efficiently prevented from inheritance, which is consistent with their localization to the GUNC (King et al., 2019; Ünal et al., 2011). We enriched for aged WT and *gim3Δ* cells by pre-staining founder cells with sulfo-NHS-LC-biotin, which permanently binds cell surface proteins, then grew cells for an average of 7 generations before isolating the biotin-marked founder cells with a streptavidin column. We first asked whether loss of Gim3 increases baseline protein aggregation. We did not observe an increase in baseline Hsp104-GFP aggregation in *gim3Δ* cells compared to WT (Figure S7I). We also examined Tub2-GFP as an additional aggregation marker and did not detect Tub2-GFP aggregation in *gim3Δ* cells (Figure S7J). Instead, Tub2-GFP signal appeared reduced on meiotic spindles, consistent with impaired tubulin incorporation into spindles, likely resulting from reduced Tub1 levels in *gim3Δ* cells. Strikingly, the Hsp104-GFP puncta observed in aged cells were inherited into spores at an elevated rate in *gim3Δ* cells compared to WT controls (Figure 5F, G). Together with the lack of increased baseline aggregation, this result suggests that Gim3 promotes the efficient exclusion of age-associated protein aggregates from gametes. Importantly, *TUB1* overexpression reduced the inheritance of Hsp104-GFP puncta in *gim3Δ* cells (Figure 5F, G), indicating that the aggregate exclusion defect is at least partly attributable to reduced Tub1 abundance downstream of Gim3.

The strong association between defects in chromosome segregation and cellular remodeling during meiosis of cells lacking Gim3 was surprising. This suggested functional mechanical linkage between the segregation of chromosomes and remodeling of other cellular compartments. To orthogonally test whether proper chromosome segregation itself is required for proper cellular remodeling, we examined cells lacking Rec8. Rec8 is a meiosis-specific cohesin subunit that is essential for sister chromatid cohesion and faithful chromosome segregation during meiosis (Klein et al., 1999). In *rec8Δ* cells, the GUNC formation measured by Nup170-GFP sequestration from chromatin was markedly reduced compared to WT (Figure 5H, I). A similar trend was also observed for Hsp104-GFP foci, phenocopying what we observed in cells lacking Gim3 (Figure 5J, K). We further found that spindle elongation was reduced in *rec8Δ* cells, particularly during meiosis II (Figure S7K). These results suggest that chromosome mis-segregation is accompanied by impaired spindle function and support a model in which spindle elongation may provide a mechanistic link between chromosome segregation and meiotic cellular remodeling. We next examined a meiosis-specific depletion allele of *SGO1* (*pCLB2-3HA-SGO1*), which impairs chromosome segregation specifically at meiosis II (Lee and Amon, 2003). In this background, both Nup170-GFP sequestration and Hsp104-GFP foci were also reduced, albeit more modestly than in *rec8Δ* cells (Figure 5H-K). Notably, chromosome mis-segregation was also less severe in *pCLB2-3HA-SGO1* cells than in *rec8Δ* cells, consistent with the milder sequestration defects observed in this background (Figure S7L). This correlation between the severity of chromosome segregation defects and impaired sequestration further supports a functional relationship between chromosome segregation and meiotic cellular remodeling. Together, these findings indicate that proper chromosome segregation supports efficient cellular remodeling and inheritance in meiosis, even beyond chromosome– and spindle-proximal structures, suggesting coupling of an unknown mechanism that enables gametes to inherit a full complement of DNA and other components.

## Discussion

In this study, we identified the prefoldin subunit Gim3 as a critical factor that safeguards chromosome segregation integrity during meiosis. Loss of Gim3 leads to severe chromosome mis-segregation and markedly reduced gamete viability. Proteomic and genetic analyses further revealed that Gim3 functions as part of the prefoldin complex to ensure the proper folding and stability of tubulin, which we found to be critical for proper spindle assembly and faithful chromosome segregation. We also discovered that Gim3 plays an important role in diverse cellular remodeling events that accompany meiotic differentiation. Moreover, induction of chromosome mis-segregation by entirely different means caused similar defects in meiotic remodeling, revealing a link between these two fundamental processes that goes beyond the coordinated gene expression program that controls both. Collectively, our results establish prefoldin as a key regulator that connects proteostasis with the maintenance of chromosomal integrity, and the coordination of chromosome segregation, cellular remodeling, and quality control during meiosis.

There are several known fundamental differences in the mechanisms of chromosome segregation in meiosis compared to mitosis that largely are related to the different chromosome behaviors during these two modes of cell division. In meiosis, there are two chromosome segregation phases following only a single DNA replication phase. Homologous chromosomes are segregated from each other at MI following their physical linkage by homologous recombination. This depends on co-orientation of sister kinetochores at MI, requiring the monopolin complex, and loss of only the cohesion complexes along chromosome arms, requiring Rec8 and Sgo1 (Nasmyth, 2002; Marston et al., 2004; Watanabe, 2012). These regulatory factors have been discovered in yeast, which has been a valuable model for uncovering conserved meiotic regulatory mechanisms. However, based on studies such as these, it is unlikely that meiotic spindles should differ fundamentally between mitosis and meiosis, as their role is similar in all cases: to separate chromosomes following attachment to kinetochores.

One surprising finding from our study is that the role of Gim3 in supporting spindle morphology and function is critical in meiosis but not mitosis. This is especially interesting given that Gim3 loss reduces Tub1 abundance to a similar degree in both cases. Why lower tubulin levels would impact meiotic but not mitotic chromosome segregation remains an open question and points to previously unrecognized differences between the mechanics of mitotic and meiotic segregation. It is possible that the demand for functionally competent tubulin is intrinsically higher during meiosis. Consistently, analysis of our previous datasets (Cheng et al., 2018) indicates that the overall protein abundance of tubulin increases during the meiotic program (Figure S8), suggesting that meiosis may be particularly sensitive to any reduction in tubulin levels. This could be based on a difference in microtubule dynamics or a feature of the formation of two spindles in rapid succession. Our finding that *spo11Δ spo13Δ* cells, which undergo only a single meiotic division, also display defective chromosome segregation in the absence of Gim3 argues against the latter model but further study is warranted in this area.

Our data suggest that short spindles alone may lead to chromosome mis-segregation. This is interesting, as previously known mutants that impact meiotic chromosome segregation largely impact either the proper regulation of linkages between chromosomes, or their attachments to the spindle. In meiotic cells lacking Gim3, we observe spindles that do not extend as far as WT cells at either MI or MII. It is possible that insufficient spatial separation of chromosomes prevents them from remaining as distinct masses. As a result, chromosomes may fail to reach nucleoplasmic regions that normally facilitate their physical segregation. Consistent with the model that low tubulin levels alone lead to short spindles and subsequent mis-segregation, we found that overexpression of a single tubulin subunit, Tub1, was sufficient to rescue the chromosome mis-segregation phenotype observed in *gim3Δ* cells. This result suggests that α-tubulin availability is selectively limiting in this context and that restoring Tub1 alone is sufficient to support functional spindle assembly. It may be that increased Tub1 abundance indirectly stabilizes the existing pool of β-tubulin by promoting efficient heterodimer formation and Tub1 levels in *gim3Δ* cells may limit the formation of polymerization-competent α/β-tubulin heterodimers. In this scenario, restoring Tub1 expression would increase the effective concentration of functional tubulin available for spindle elongation without requiring a concomitant increase in Tub2 synthesis.

The observation that induction of chromosome mis-segregation itself causes defects in cellular remodeling suggests that these processes are not merely parallel but hierarchically linked. These defects in remodeling, which include sequestration of NPC components away from chromatin and removal of Hsp104-associated protein aggregates from aged cells reflect mechanistic linkage between the distribution of chromosomes and other cellular components. The improper segregation of chromosomes appears to affect the ability of Hsp104-marked protein aggregates to be sequestered and removed during meiosis. It also causes improper sequestration of NPC components and reduces the ability of meiotic cells to efficiently form gametes. These two observations have been previously linked in mutants that impair gamete membrane formation (King et al., 2019). Whether this is the root cause of the defect in Gim3-deficient cells is not yet known. However, it is clear that unexpected connections exist between multiple aspects of cellular remodeling and inheritance. Determining the basis of these linkages is an exciting future direction. One possibility is that these remodeling events depend on microtubule-based organization within meiotic cells. Microtubules are known to mediate the transport and sequestration of protein aggregates and other cellular components (Johnston et al., 1998; Egan et al., 2015). Thus, the reduced Tub1 levels caused by loss of Gim3 may impair microtubule-dependent organization of intracellular components, which could contribute to the remodeling defects observed in meiosis. Also, given the tight packing of components inside cells during the dramatic morphological changes that accompany gametogenesis, the forces associated with chromosome segregation may drive changes to other cellular compartments.

Or perhaps completion of chromosome segregation may provide a structural or signaling cue that triggers subsequent remodeling events, such as nuclear envelope reorganization and selective degradation of nuclear pore components. In this view, segregation errors could interfere with this signaling cascade, thereby impairing the proper reorganization and compartmentalization of cellular structures. From a physiological standpoint, this coupling could ensure that genome integrity and cellular rejuvenation are achieved simultaneously, thereby safeguarding the transmission of genetic information while excluding the unnecessary cellular components to the next generation. Functionally, our results indicate that prefoldin function, and resultant tubulin stability, play a critical role in linking microtubule integrity and cellular reorganization to secure the fidelity and quality of gametogenesis.

Notably, although both nuclear pore component (Nup170-GFP) and protein aggregate (Hsp104-GFP) behavior are misregulated in cells with chromosome segregation defects, their apparent fates differ. Nup170-GFP masses are not inherited and are instead degraded together with associated chromosome masses in spores that fail to cellularize, whereas Hsp104-GFP puncta are aberrantly inherited into gametes when segregation is impaired. This divergence is intriguing and suggests that different classes of cellular components are subject to distinct regulatory mechanism during gametogenesis, even when their initial mislocalization arises from a common defect in chromosome segregation. Recent studies have similarly reported differential behaviors of nuclear pore components and protein aggregation during gamete formation (Ruediger et al., 2025), supporting the idea that these processes are not uniformly governed by passive co-segregation with chromosomes. Instead, spores may actively discriminate between cellular components, selectively directing them toward degradation or inheritance. Such selectivity could allow gametes to fine-tune their proteome and organelle composition, rather than simply inheriting or excluding components based solely on spatial proximity.

Mechanisms underlying meiosis are conserved from yeast to mammals, suggesting that the findings of this study may have broader implications beyond budding yeast. Indeed, several lines of evidence point to a potential role of prefoldin subunits in mammalian gametogenesis. In humans, a study examining chromatin states and sperm quality reported that CpG hypomethylation of PFDN1, PFDN4, PFDN5, and PFDN6 correlates with reduced sperm motility (Figure S9A; Pacheco et al., 2011). Furthermore, transcriptome analyses of sperm from patients with teratozoospermia revealed decreased expression of PFDN1, PFDN4, and PFDN5 (Figure S9B, Platts et al., 2007), while analyses of human spermatogenic arrest showed altered expression of PFDN2 and CCT4, which we identified as Gim3 interactors during meiosis (Figure S9C, Kui et al., 2019). Importantly, genetic studies in mice and Drosophila demonstrated that loss of prefoldin leads to abnormal spermatogenesis, further supporting the conserved requirement of prefoldin in gamete development (Lee et al., 2011; Delgehyr et al., 2012). However, whether these abnormalities arise directly from errors in meiotic chromosome segregation or cellular remodeling remains unknown. Given that our current work in yeast establishes a direct link between prefoldin function in supporting tubulin levels and meiotic chromosome segregation fidelity, it would be intriguing to investigate whether similar mechanisms underlie meiosis in mammalian systems.

## Limitations of the study

Although our findings uncover an unexpected link between the prefoldin complex and meiotic remodeling, this study was conducted in a single organismal context. Thus, whether the coupling between protein folding capacity and chromosome segregation represents a conserved principle across eukaryotes remains speculative. In assaying chromosome segregation, we assessed only chromosome V specifically, along with overall chromatin segregation. Whether differences in the degree of mis-segregation of different chromosomes is seen in cells lacking Gim3 is unknown. Finally, the functional significance of prefoldin-mediated cytoskeletal regulation during stress or aging, which may also involve cytoskeletal vulnerabilities, has yet to be explored. Expanding these analyses to other model systems or mammalian germ cells will be essential to assess the broad relevance of our findings.

## Methods

### Strain construction

Strains were constructed in the SK1 background of *Saccharomyces cerevisiae*. Strains and plasmids used for this study are listed in Supplementary file 5. Deletion and C-terminal tagging at endogenous loci were performed using previously described PCR-based methods unless otherwise specified (Janke et al., 2004; Longtine et al., 1998; Powers et al., 2022; Sheff and Thorn, 2004).

The following strains were constructed in a previous paper: Htb1-mCherry (Matos et al., 2008), CenV GFP (Lee and Amon, 2003), GFP-Tub1 (Falk et al., 2016), Spc42-mCherry (Miller et al., 2012), *spo11Δ*, *spo13Δ* (Lee and Amon, 2003), Nup170-GFP (King et al., 2019), *flo8Δ* (Boselli et al., 2009), Hsp104-GFP (Ünal et al., 2011), Cit1-GFP (Sawyer et al., 2019), HDEL-GFP (Otto et al., 2021), *tpk1-as tpk2Δ tpk3Δ* (Stephan et al., 2009), *mad2Δ* (Sheltzer et al., 2011) and *rec8Δ* (Klein et al., 1999).

### Yeast growth and sporulation conditions

Mitotic cells were grown in YPD (1% yeast extract, 2% peptone, 2% glucose, 22.4 mg/L uracil, and 80 mg/L tryptophan) at 30 °C, with exponentially growing cells grown from an OD_600_ of 0.2. For meiotic time courses, strains were inoculated in YPD for 24 hours, then diluted to an OD_600_ of 0.2 in BYTA (1% yeast extract, 2% bacto tryptone, 1% potassium acetate, and 50 mM potassium phthalate) and grown for 16 hours at 30 °C. After reaching an OD_600_ ≥5, the cells were pelleted, washed in sterile MilliQ water, and resuspended in SPO to OD_600_ = 1.85. SPO was 0.5% potassium acetate alone, 1% potassium acetate alone, or 2% potassium acetate supplemented with amino acids (40 mg/L adenine, 40 mg/L uracil, 10 mg/L histidine, 10 mg/L leucine and 10 mg/L tryptophan); the media’s pH was adjusted to 7 with acetic acid. Meiotic cultures were shaken at 30 °C for the duration of the experiment. At all stages, the flask size was 10 times the culture volume to ensure proper aeration.

### Sporulation conditions for *tpk1-as*

Cells were grown overnight in YPD, diluted to BYTA for 16 hours, and subsequently shifted to YPD (OD_600_ = 1.0). Rapamycin was added to cells at a final concentration of 1000 ng/mL. 1NM-PP1 was added to cells at a final concentration of 3 μM.

### CRISPR screening

#### • Library Design

CRISPR screening was conducted using CRISPEY, a Cas9 and retron-based method for parallel and precise genome editing (Sharon et al., 2018). Libraries of donor-guide oligonucleotides were designed to systematically perturb sORFs and tORFs in Saccharomyces cerevisiae. The design strategy was implemented using custom scripts provided by Helen Sakharova (https://github.com/hsakharova/library_design). All oligonucleotide sequences used in this study are provided in Supplementary File 1. Each oligonucleotide insert was 194 bp in length and consisted of a 20-bp 5’ homology arm, a 100-bp donor sequence, a 34-bp retron scaffold, a 20-bp guide sequence, and a 20-bp 3’ homology arm. These inserts were designed for cloning into the pZS165 plasmid backbone using NotI-HF digestion and NEBuilder HiFi DNA Assembly (New England Biolabs), followed by transformation into E. coli for library preparation (Sharon et al., 2018).

For each sORF, we designed donor-guide oligonucleotides that would disrupt the start codon by deleting the second base (ATG to AG). In sORFs with downstream alternative start codons, or if no guide could be found to target the start codon, we also designed donor-guide oligonucleotides that would introduce early (< 20% ORF length) stop codons along with a frameshift (NNNN to TAA). We prioritized guides that would minimize the number of downstream alternative start codons. If any downstream start codons remained, we designed donor-guide oligonucleotides that would cause a mid-ORF (< 60% ORF length) stop codon and frameshift.

For tORFs, we designed donor-guides to disrupt the putative start codon of the truncation with a MET to ILE mutation. In order to help ensure that any change was caused by the removal of the start codon rather than the introduction of ILE, we also included MET to ALA mutations. Frameshifts were avoided in order to preserve the production of the longer isoform. If no guide targeting the putative start codon could be found, the next start codon downstream of the putative start codon was targeted instead.

Where possible, we included multiple different donor-guide oligonucleotides targeting the same mutation. Guides were filtered to have no off targets with fewer than three mismatches to the guide. Furthermore, in order to prevent repeat cutting of the genomic DNA after editing, we included only guide-donors that would create mismatches with the seed region of the guide (the 7 bases upstream of the PAM). For desired mutations that could not be targeted under these strict restrictions, we included guides that did not fulfill them, as long as all potential off-target sites had at least one mismatch with the seed region of the guide. However, we considered these guides as less reliable, and included additional guide-donors targeting locations further downstream.

As positive controls, we included guide-donor oligonucleotides that would disrupt ∼50 annotated genes with known mitotic or meiotic phenotypes. We also included 30 random guide sequences with no predicted on-target sites in the yeast genome as negative controls.

#### • Yeast Library Preparation

Library oligonucleotides were PCR-amplified with pUB14226/14227 (Supplementary file 5). Amplified fragments were purified and assembled into NotI-HF (New England Biolabs) – digested pUB2965 vector using NEBuilder HiFi DNA Assembly (New England Biolabs). Ligation products were ethanol-precipitated and electroporated into E. coli (MegaX). Colonies were expanded on large LB agar plates to achieve >30× library coverage. Plasmids were recovered by midiprep (Qiagen) and digested with NotI-HF and CIP (New England Biolabs) to remove empty vector before yeast transformation

Yeast strains were cultured in YPD to OD_600_ ≈ 0.6, harvested, and washed with ice-cold water and 1 M sorbitol. Cell pellets were resuspended in freshly prepared LiTE buffer (10 mM Tris-HCl pH 7.9, 1 mM EDTA, 100 mM LiAc, 25 mM DTT (dithiothreitol)), followed by a final wash with sorbitol. Cells were resuspended in 1 M sorbitol (100-200 µL per pellet). For electroporation, 5 µg plasmid DNA per 100 µL cells was added, and 100 µL aliquots were transferred to 0.2 cm cuvettes. Electroporation was performed at 1.5 kV, 25 µF, 100 Ω. Immediately after pulsing, 1 mL of pre-warmed YPD was added for recovery. Cells were transferred to selective YPD + G418 and incubated at 30 °C with shaking. Colonies were pooled after overnight recovery, washed, and stored in 15% glycerol at ∼50 OD/mL at –80 °C.

For library editing, a fresh aliquot of the desired yeast cDNA library was thawed and used to inoculate a YPR (1% yeast extract, 2% peptone, 2% raffinose, 22.4 mg/L uracil, and 80 mg/L tryptophan) + 320 µg/mL G-418. The yeast library was recovered at 30 °C overnight and then cells were washed and resuspended with YPGR media (1% yeast extract, 2% peptone, 2% raffinose, 2% galactose, 22.4 mg/L uracil, and 80 mg/L tryptophan) + 320 µg/mL G-418for 48 hours. The edited yeast library was stored in 15% glycerol at –80 °C.

#### • Enrichment of spores

To enrich for sporulated cells, the edited yeast library was inoculated into BYTA and then SPO and cultured for 12 h prior to harvest. Cultures were washed once in sterile water and once in 1 M sorbitol. Cell pellets were resuspended in 5 mL SCE buffer (1 M sorbitol, 0.1 M sodium citrate pH 5.8, 10 mM EDTA) supplemented with 50 µL 1 M DTT and zymolyase to a final concentration of 1 mg/mL and incubated for 15 min at 30 °C with shaking. Following treatment, SDS was added to a final concentration of 1% and incubated for 5 min to lyse unsporulated cells. Efficient removal of mitotic cells was confirmed microscopically. The remaining spores were washed in water, resuspended in 15% glycerol, and stored at –80 °C

#### • Plasmid extraction

For plasmid recovery, yeast stocks were thawed from –80 °C and pelleted at 2000 rcf for 2 min. Pellets were resuspended in P1 buffer (Qiagen) supplemented with 2.5 µL 1 M DTT and 25 µL of 10 mg/mL zymolyase, followed by incubation for 15 min at 37 °C. Cells were disrupted by bead beating (3 min), and plasmids were isolated using a QIAprep spin column kit (Qiagen) with additional washes in PB and PE buffers (Qiagen). DNA was eluted twice in 15 µL H₂O, and plasmid concentration was measured by Qubit fluorometry (Thermo Fisher).

#### • Library sequencing

Plasmids recovered from yeast were used as templates for two-step PCR to generate Illumina-compatible sequencing libraries. In the first PCR, primers containing staggered overhangs (pUB11732/12227-12234, Supplementary file 5) were used to amplify the donor-guide insert. The second PCR appended Illumina adapters and indices using primers oCJC161 and oCJC65/66/72/73 (Supplementary file 5). PCR products of the expected size were purified by gel extraction and quantified using TapeStation analysis (Agilent). Libraries were sequenced on an Illumina platform.

#### • Data analysis

For each donor-guide construct, the 5’ terminal 60 bp of the donor sequence was extracted and used as the reference for mapping. Processed reads were aligned and quantified using sgcount (https://noamteyssier.github.io/sgcount/), and the resulting count tables were analyzed with MAGeCK (Li et al., 2014) to assess differential enrichment or depletion of guides across conditions

### Live-cell imaging

Images were acquired using a DeltaVision Elite wide-field fluorescence microscope (GE Healthcare) and a PCO Edge scientific complementary metal-oxide-semiconductor camera, with softWoRx software and a 60× NA1.42 oil-immersion Plan Apochromat objective. Live-cell imaging was performed exactly as described in (King et al., 2019). In short, cells were imaged in an environmental chamber heated to 30 °C, using concanavalin A-coated glass-bottom 96-well plates. Cultures of meiotic cells in SPO were adhered to wells, and 100 µL of SPO was added to each well. Images were deconvolved using softWoRx software (GE Healthcare) using 3D iterative constrained deconvolution with 15 iterations and enhanced ratio.

### Spore viability assay

To assess spore viability, 100 µL aliquots of SPO cultures collected after 24 h at 30 °C were spun in 1.5 mL tubes at 1,900 rcf for 2 min at room temperature and SPO was removed from cell pellets by pipetting. To digest asci and allow for manual separation of individual spores on agar plates, cells were resuspended in 20 µL of (1 mg/mL) 100T zymolyase (cat. no. 08320932, MP Biomedicals or cat. no. 320932; VWR) and incubated at room temperature for 6-7 min. To stop digestion, 180 µL of MilliQ H_2_O was added to each sample, and 20 µL of digested cells were then pipetted down the midline of YPD agar plates (1X YEP with 2% dextrose). Isolation of spores from the same tetrad was performed by microdissection using a fiberoptic needle attached to a Zeiss light microscope.

### Immunopurification

Yeast cultures were induced to enter meiosis by incubation in SPO at 30 °C. At indicated timepoints, 50 mL of culture (∼92.5 OD units) or an equivalent amount of mitotic culture was harvested. Cells were collected into 50 mL tubes on ice, and PMSF was added to a final concentration of 2 mM. After brief mixing, cells were pelleted at 3,000 rpm for 2 min at 4 °C. Supernatant was removed, and the pellet was resuspended in 1 mL of supernatant in a screw-cap tube. After centrifugation at 15,000 rpm for 1 min at 4 °C, the supernatant was discarded, and the pellet was snap-frozen in liquid nitrogen and stored at –80 °C.

Pellets were thawed on ice, and zirconia beads (250 µL) were added. Lysis was performed by bead beating (FastPrep, 6.0 m/s, 2 × 20 s, with 2 min cooling on ice between runs) in 250 µL of NP-40 lysis buffer supplemented with protease inhibitors (150 mM Tris-HCl (pH 7.5), 150 mM NaCl, 1% NP-40, and 5% glycerol, supplemented with 0.5 mM MgCl₂, PhosSTOP phosphatase inhibitor (Roche; 1 tablet per 10 mL buffer), cOmplete protease inhibitor (Roche; ¼ tablet per 10 mL buffer), 0.1 mg/mL AEBSF (Sigma), and 2.7 µM Pepstatin A (Sigma)). Lysates were clarified by centrifugation at 8,000 g for 20 s at 4 °C, and supernatants were transferred to 1.5 mL tubes. After a second centrifugation at 21,000 g for 20 min at 4 °C, cleared lysates were snap-frozen in liquid nitrogen and stored at –80 °C.

### Sample preparation for the LC-MS/MS analysis

#### Total Lysate

Clarified lysates prepared in the ribosome profiling protocol were subjected to downstream MS analysis. Proteins were precipitated and desalted using the SP3 method, as described in (Hughes et al., 2019). The protein concentration was determined by using the Pierce™ BCA protein assay kit (Thermo Scientific) based on the manufacturer’s instructions. 50 µg from each lysate were processed further. Disulfide bonds were reduced with 5 mM dithiothreitol 45 min at 600 rpm and 25°C and cysteines were subsequently alkylated with 10 mM iodoacetamide for 45 min in the dark at 600 rpm and 25°C. SDS was added to each sample for a final concentration of 1%. Proteins were then precipitated on 10 µL (concentration 50 mg/mL) speedBead magnetic carboxylated modified beads (total of 500 1:1 mix of hydrophobic and hydrophilic beads, cat# 6515215050250, 45152105050250, GE) by addition of 100% ethanol in a 1:1 (vol:vol) sample:ethanol ratio followed by 15 min incubation at 25°C, 1000 rpm. Protein-bound beads were washed in 180 µL 80% ethanol 3 times and proteins were digested off the beads by addition of 1 µg sequencing grade modified trypsin (Promega) in 100 mM Ammonium bicarbonate, incubated 16 hrs at 25°C, 600 rpm. Beads were removed and the resulting tryptic peptides evaporated to dryness in a vacuum concentrator. Dried peptides were then reconstituted in 60 μL of 3% acetonitrile / 0.2% Formic acid for LC-MS/MS.

#### IP samples

130 μL of 50 mM Tris/HCl (pH 8) was added to each sample protein eluate and disulfide bonds were reduced with 5 mM dithiothreitol (DTT) for 45 min at 600 rpm and 25°C. Cysteines were subsequently alkylated with 10 mM iodoacetamide (IAA) for 45 min in the dark at 600 rpm and 25°C. Proteins were then digested by adding 0.5 μg sequencing grade modified trypsin (Promega) for 16 h at 600 rpm and 25°C. After digestion, samples were acidified with a final concentration of 1% formic acid. Tryptic peptides were desalted on C18 StageTips according to (Rappsilber et al., 2007), dried in a vacuum concentrator, and reconstituted in 13 μL of 3% acetonitrile / 0.2% formic acid for LC-MS/MS.

### LC-MS/MS analysis

#### Total Lysate

Approximately 1 μg of total peptides were analyzed on a Waters M-Class UPLC using a 15 cm x 75 µm IonOpticks C18 1.7 µm column coupled to a benchtop Thermo Fisher Scientific Orbitrap Q Exactive HF mass spectrometer. Peptides were separated at a 400 nL/min flow rate with a 150-minute gradient, including sample loading and column equilibration times. Data were acquired in data-independent mode using Xcalibur software (4.5.474.0). MS1 spectra were measured with a resolution of 120,000, an AGC target of 3e6, and a scan range from 350 to 1600 m/z. 35 isolation windows of 36 m/z were measured at a resolution of 30,000, an AGC target of 3e6, normalized collision energies of 22.5, 25, 27.5, and a fixed first mass of 200 m/z. Proteomics raw data were analyzed with Spectronaut software version 20.0.250606.92449 (Bruderer et al., 2015) using the yeast UniProt database (UP000002311) under the default directDIA analysis setting with cross-run normalization and global imputation.

#### IP samples

LC-MS/MS analysis was performed on a Q-Exactive HF. Around 0.5 μg of total peptides were analyzed on a Waters M-Class UPLC using a 15cm Ion-Optics column (1.7um, C18, 75um x 15cm) coupled to a benchtop ThermoFisher Scientific Orbitrap Q Exactive HF mass spectrometer. Peptides were separated at a flow rate of 400 nL/min with a 90 min gradient, including sample loading and column equilibration times. Data was acquired in data-dependent mode. MS1 spectra were measured with a resolution of 120,000, an AGC target of 3e6 and a mass range from 300 to 1800 m/z. MS2 spectra were measured with a resolution of 15,000, an AGC target of 1e5 and a mass range from 200 to 2000 m/z. MS2 isolation windows of 1.6 m/z were measured with a normalized collision energy of 25.

Proteomics raw data was analyzed by MaxQuant (v2.0.3.0) (Cox and Mann, 2008) using a UniProt yeast database (UP000002311), and MS/MS searches were performed under default settings with iBAQ and LFQ quantification and match between the runs. The MS1 iBAQ intensities were normalized such that each sample intensity values added up to exactly 1,000,000, therefore each protein group value can be regarded as a normalized microshare. A pseudocount of 2 was added to each of these normalized iBAQ microshare values and then these values were log2 transformed.

### Western blotting

Protein samples were prepared by TCA (trichloroacetic acid) treatment of cells. For meiotic samples, 1.8 mL culture was mixed with 200 µL of 50% TCA (5% final concentration) and incubated at 4 °C for 12-24 h. For mitotic samples, 3.42 OD units of culture was spun down for 2 min at 3,000 rcf and resuspended in 5% TCA, and incubated at 4 °C for 12-24 h. All samples were precipitated for 5 min at 20,000 rcf and washed in 1 mL acetone. The acetone was aspirated, and samples were allowed to dry for ≥20 min. Pellets were resuspended by bead beating for 5 min in 100 µL Tris-EDTA buffer (10 mM Tris-HCL, 1 mM EDTA, pH 8) supplemented with 3 mM DTT and 1× protease inhibitors (Roche) with 100 µL acid-washed glass beads. 50 µL of 3× SDS loading buffer was added, and samples were incubated at 95 °C for 5 min and spun down for 5 min at 20,000 rcf. 4 µL sample was loaded onto a Bis-Tris acrylamide gel, separated at 150 V for 50 min, and transferred to a nitrocellulose membrane using the TransBlot Turbo system (Bio-Rad). Blots were blocked and probed overnight at 4 °C with one or more of the following antibodies: anti-hexokinase (RRID:AB_219918, 100-4159, 1:15,000; Rockland), anti-Tub1 (RRID:AB_325003, MCA78G, 1:1,000; BioRad), and anti-V5 (RRID:AB_2556564, R960-25, 1:2,000; Thermo Fisher Scientific). Blots were washed in PBS-T and incubated for 2 h in IRDye secondary antibodies (RRID:AB_10956166, 926-68071, 1:20,000; LI-COR, RRID:AB_2721932, 925-32219 1:20,000; LI-COR and RRID:AB_621847, 926-32212, 1:20,000; LI-COR). Blots were imaged and quantified using the Odyssey system (LI-COR).

### Ribosome profiling

Ribosome profiling was performed as in (Hollerer et al., 2021; Powers and Brar, 2021). Briefly, cells were treated with cycloheximide for 30s then filtered and flash frozen. Extracts were milled under cryogenic temperatures and stored at –80 °C in aliquots. RNA extracted from monosomes was extracted and fragments ∼28-32nt were collected. Libraries were prepped using linker ligation and rRNA fragments were depleted from samples using biotinylated anti rRNA oligos. Samples were sequenced using 50nt single end reads on a HS4000 or using 50nt single end reads on a NovaSeq 6000. Matched mRNA-seq libraries were prepared with the same library prep protocol with the following changes. Poly A selection was used to isolate mRNA from extracted total RNA samples which were subsequently fragmented and libraries were created with fragments ∼35-80nt. No rRNA depletion was performed on these samples.

Sequencing reads were processed using a custom Snakemake pipeline. Adaptor sequences were trimmed with fastx_clipper, and reads aligning to rRNA were removed with bowtie2 (Langmead and Salzberg, 2012). Remaining reads were aligned to the Saccharomyces cerevisiae SK1 reference genome using STAR (Dobin et al., 2013). Only uniquely mapping reads (MAPQ ≥ 50) were retained, followed by sorting and indexing with samtools (Li et al., 2009). Read periodicity and A-site offsets were determined with fp-framing, and gene-level ribosome occupancy was quantified using fp-count. Strand-specific wiggle files were generated with wiggle-track for downstream visualization. Visualization was performed using Integrative Genomics Viewer (Robinson, 2011).

### Aged cell enrichment

Aged cells were enriched using a biotin-labeling and magnetic-sorting assay (Smeal et al., 1996). Cells were grown in YPD at room temperature or 30 °C overnight until saturation (OD_600_ ≥10) and then diluted to a cell density of OD_600_ = 0.2 in a new YPD culture. Cells were harvested before the cultures reached OD_600_ = 1 and were labeled with 8 mg/mL EZ-Link Sulfo-NHS-LC-biotin (ThermoFisher Scientific) for 30 min at 4 °C. Biotinylated cells were grown for 12-16 hours in YPD with 100 μg/mL ampicillin at 30 °C. Cells were subsequently harvested and mixed with 100 µL of anti-biotin magnetic beads (Miltenyi Biotechnology) for 15 min at 4 °C. Cells were washed with PBS pH 7.4, 0.5% BSA (bovine serum albumin) buffer and sorted magnetically using LS depletion columns with a QuadroMacs sorter following the manufacturer’s protocol. Aged cells were subsequently washed once with SPO. The cell mixture was resuspended with SPO at a cell density of OD_600_ = 1.85 with 100 μg/mL ampicillin and incubated at 30 °C.

### Fluorescence microscopy

Images were acquired using a DeltaVision Elite wide-field fluorescence microscope (GE Healthcare). Images were generated using a 60x/1.42 NA oil-immersion objective lens. Images were deconvolved using softWoRx imaging software (GE Healthcare). Unless otherwise noted, images were maximum intensity z-projected over the range of acquisition in FIJI (Schindelin et al., 2012).

### Image quantification

Meiosis progression (Figure S1C) was assessed by DAPI staining. Cells that had not yet initiated meiosis I and displayed a single DAPI-stained nuclear mass were defined as mononucleated cells. Because variability between experimental batches was observed, values were normalized within each experiment.

Chromatin segregation was assessed using Htb1-mCherry fluorescence. Chromatin mis-segregation events were manually scored during nuclear division when the chromatin signal remained stretched between separating nuclei or when unequal chromatin masses were observed.

Spindle structures were analyzed using GFP-Tub1 and Spc42-mCherry signals. In Fig. 3F and Fig. 3G, cells displaying clearly elongated spindle structures were manually counted. For Figure S4, GFP-Tub1 fluorescence intensity located between Htb1-mCherry signals was quantified using FIJI (Schindelin et al., 2012). Spindle length measurements in Figure 3 were performed by measuring the distance between Spc42-mCherry foci using FIJI (Schindelin et al., 2012). During meiosis I, the distance between the two Spc42-mCherry foci was measured. During meiosis II, the longest distance among the four Spc42-mCherry foci was used as the spindle length.

Localization of Nup170-GFP and Hsp104-GFP was analyzed by manual scoring. As previously reported (King et al., 2019), these proteins are normally segregated away from chromatin and sequestered into the Gametogenesis-Uninherited Nuclear Compartment. Cells were scored for aggregate formation and sequestration from chromatin. Cells were considered defective when sequestration was incomplete or when GFP signals overlapped with Htb1-mCherry.

### Benomyl treatment

Benomyl medium was prepared as described by (Hochwagen et al., 2005). Briefly, a 30 mg/mL benomyl stock solution (Methyl 1-(butylcarbamoyl)-2-benzimidazolecarbamate; Aldrich, 95%) was freshly prepared in DMSO on the day of the experiment. The stock was added to boiling SPO to a final concentration of 20 µg/mL without swirling, followed by gentle mixing until the solution was completely clear with no visible flakes. The medium was then cooled slowly to 30 °C and used the same day. Meiotic cells were transferred from regular sporulation medium to benomyl medium by filtration.

### Statistical analysis

The statistical tests used for each dataset are indicated in the corresponding figure legends. Statistical analyses were performed using GraphPad Prism version 10. Welch’s t-test was used for two-group comparisons. Data distributions for t-test analyses were assumed to be normal but were not formally tested. Mann-Whitney U tests were used for two-group comparisons when the data were analyzed using a nonparametric test. For comparisons involving three or more groups, one-way analysis of variance (ANOVA) followed by Dunnett’s test or Tukey’s multiple comparison test was applied.

## Data availability

- This paper does not report original code.
- Raw ribosome profiling data are deposited at NCBI GEO with accession numbers GSE314037.
- Any additional information required to reanalyze the data reported in this paper is available from the lead contact upon request.

## Acknowledgments

We thank Michael Rape and members of the Brar and Ünal labs for critical feedback on the manuscript. Sequencing was performed at the UCSF CAT, supported by UCSF PBBR, RRP IMIA, and NIH 1S10OD028511-01 grants. This work was supported by the National Institutes of Health funding to G.A.B (R35GM134886), M.J. (R35GM152258) and E.Ü. (R01AG071801). E.Ü. was also funded by the Astera Institute. B.Styler was funded by NIH training grant (T32 GM007232) and an NSF predoctoral fellowship (DGE 2146752). N.K. was funded by an overseas research fellowship and research fellowship for young scientists from Japan Society for the Promotion of Sciences.

## Author Contributions

N.K.: conception and design, collection and assembly of data, data analysis and interpretation, and manuscript writing. B. Stekovic and M.J.: collection, analysis, and interpretation of MS experiments. H.S. and L.L.: design of the screening library and assistance with CRISPEY implementation. B. Styler and E.Ü.: conception and data interpretation for cellular remodeling experiments. G.A.B.: study conception and design, data interpretation, financial and administrative support, supervision, and manuscript writing. All authors: revision and final approval of the manuscript.

## Use of AI tools

During the preparation of this work, N.K. used ChatGPT and DeepL in order to improve the English text. After using these tools, the authors reviewed and edited the content as needed and take full responsibility for the content of the published article.

## Declaration of Interests

The authors declare no competing interests.

## Supplementary Figure Legends

**Supplementary Figure 1.**
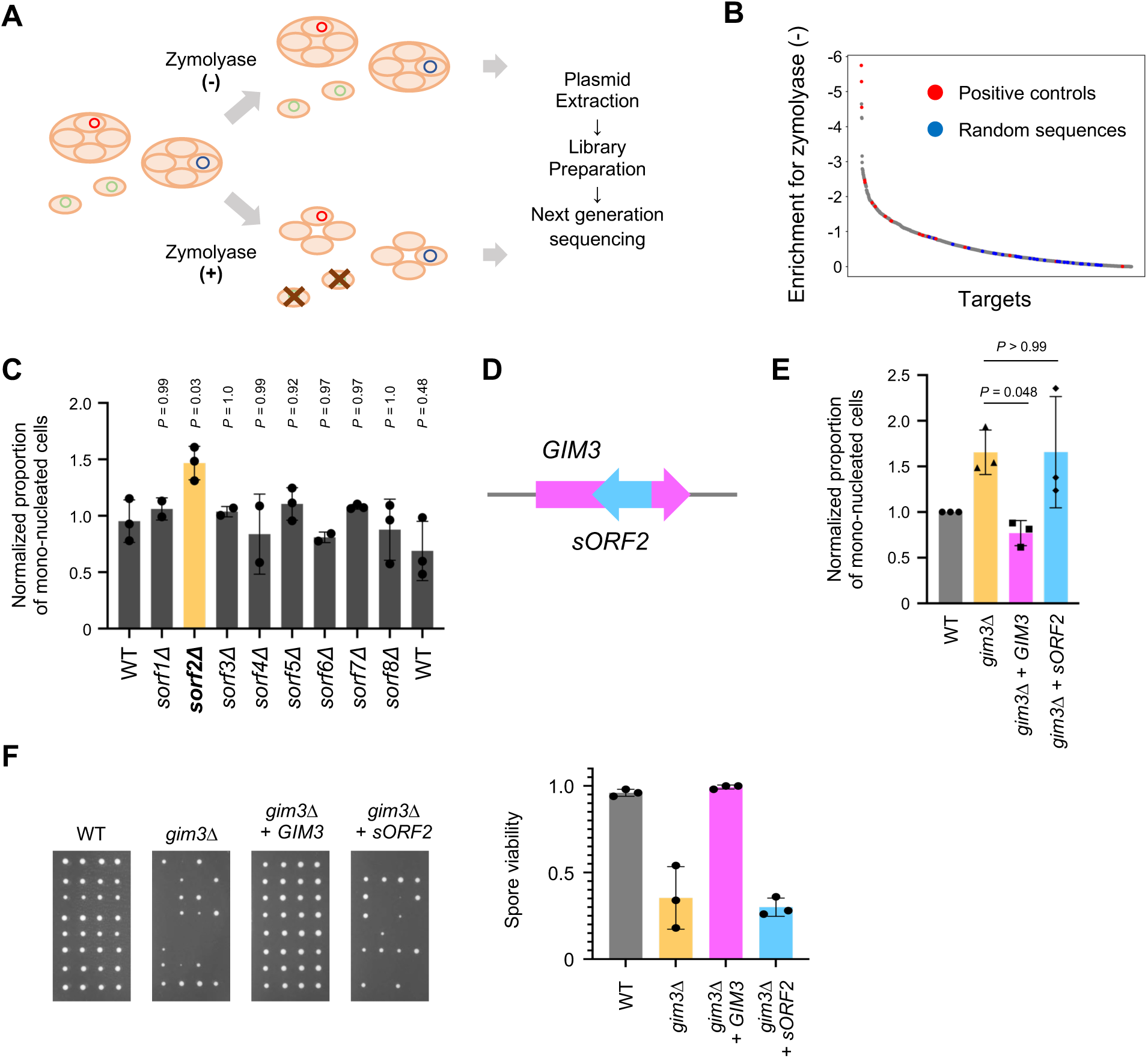
CRISPR screening and following experiments identify Gim3 as meiotic regulator. (A) Screening scheme. A CRISPEY library targeting short open reading frames (sORFs) and truncated ORFs (tORFs) was constructed and introduced into yeast. After meiosis induction, spores were enriched by zymolyase treatment, followed by plasmid extraction, library preparation, and next-generation sequencing. (B) Enrichment scores for resistance to zymolyase treatment. Random sequences are shown in blue and positive control sequences are shown in red. (C) Validation of top sORF hits identified in (B). Deletion strains for the indicated sORFs were constructed, and meiotic progression was assessed by DAPI staining at 6 hours in SPO. The y-axis represents the normalized proportion of mono-nucleated cells. N = 2-3; Dunnett’s test. More than 100 cells were quantified per replicate. (D) Genomic locus of *GIM3* and *sORF2*. *sORF2* resides within the *GIM3* locus and is transcribed in the reverse direction. (E) Complementation assay. *gim3Δ* strains were transformed with plasmids expressing either *GIM3* or *sORF2*, and meiotic progression was assessed by DAPI staining. The y-axis represents the normalized proportion of mononucleated cells at 6 hours in SPO. N = 3. Dunnett’s test. More than 100 cells were quantified per replicate. (F) Spore colony growth of WT, *gim3Δ* with empty cassette, and *gim3Δ* with *GIM3* or *sORF2* complementation after 2 days at 30 °C (left). Quantification of spore viability (right). N = 3; data are represented as mean ± SD. Dunnett’s test. 64-80 spores were quantified per replicate. The results for WT*, gim3Δ* carrying the empty cassette, and *gim3Δ* complemented with *GIM3* are identical to those shown in Figure 2C, D.

**Supplementary Figure 2.**
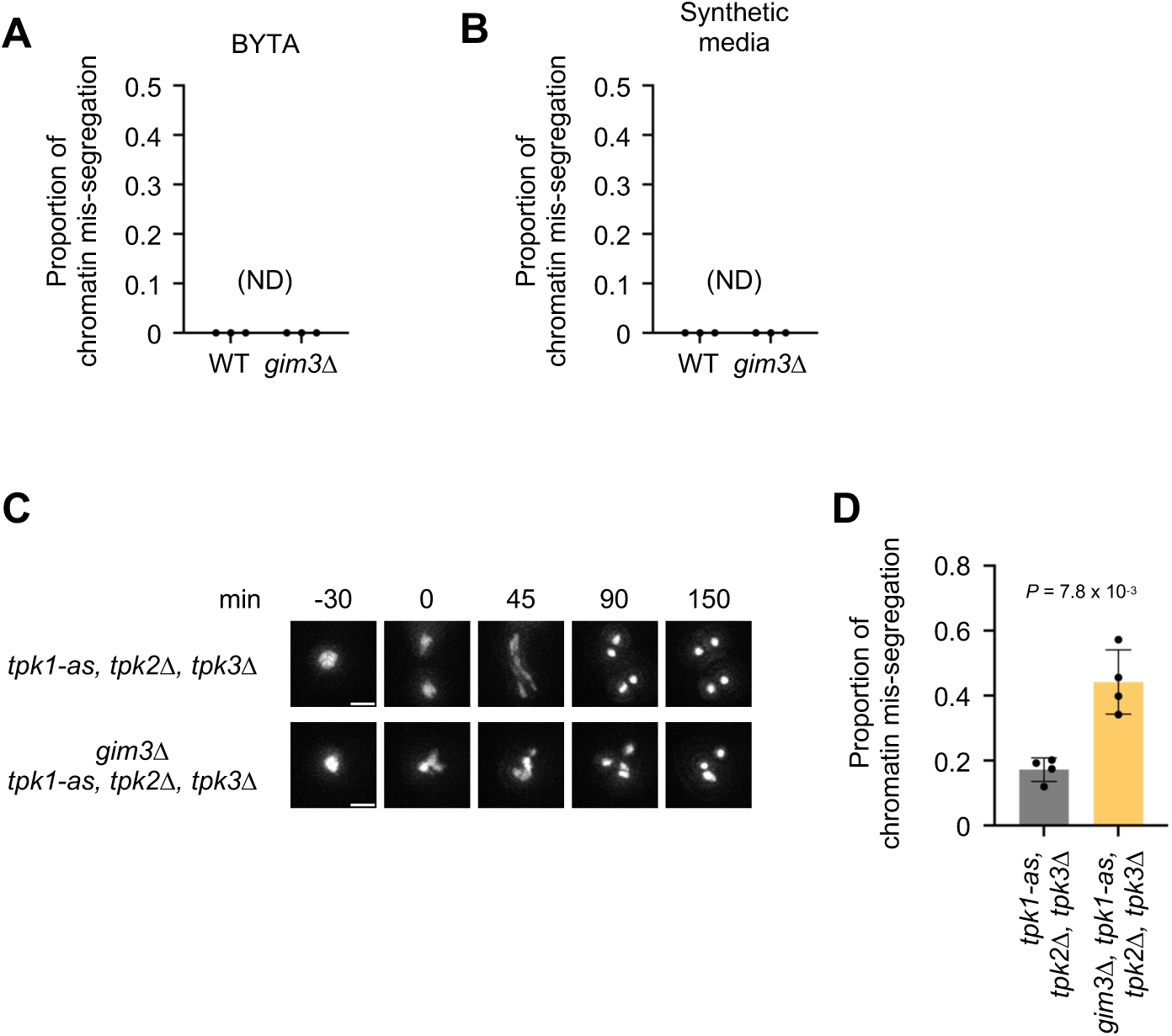
Chromatin mis-segregation was not detected during mitosis in *gim3Δ* cells grown in pre-sporulation or synthetic media. (A) Quantification of chromatin mis-segregation during mitosis in WT and *gim3Δ* cells in BYTA. Htb1-mCherry imaging was performed every 10 min during mitosis. N = 3; ND, not detected; 71-80 cells were quantified per replicate. (B) Quantification of chromatin mis-segregation during mitosis in WT and *gim3Δ* cells in synthetic media. Htb1-mCherry imaging was performed every 15 min during mitosis. N = 3; ND, not detected; 27-55 cells were quantified per replicate. (C) Live-cell imaging of Htb1-mCherry in *tpk1-as tpk2Δ tpk3Δ* cells (top) and *gim3Δ tpk1-as tpk2Δ tpk3Δ* cells (bottom) in YPD treated with 1NM-PP1 and rapamycin. Cells were imaged every 15 min during meiosis. Yellow arrowheads indicate chromatin mis-segregation. Time is shown relative to the onset of anaphase I (0 min). Scale bars, 3 µm (D) Quantification of Htb1-mCherry mis-segregation events during MI (left) and MI&MII (right). N = 4; data are represented as mean ± SD; t-test. 100-130 cells were quantified per replicate.

**Supplementary Figure 3.**
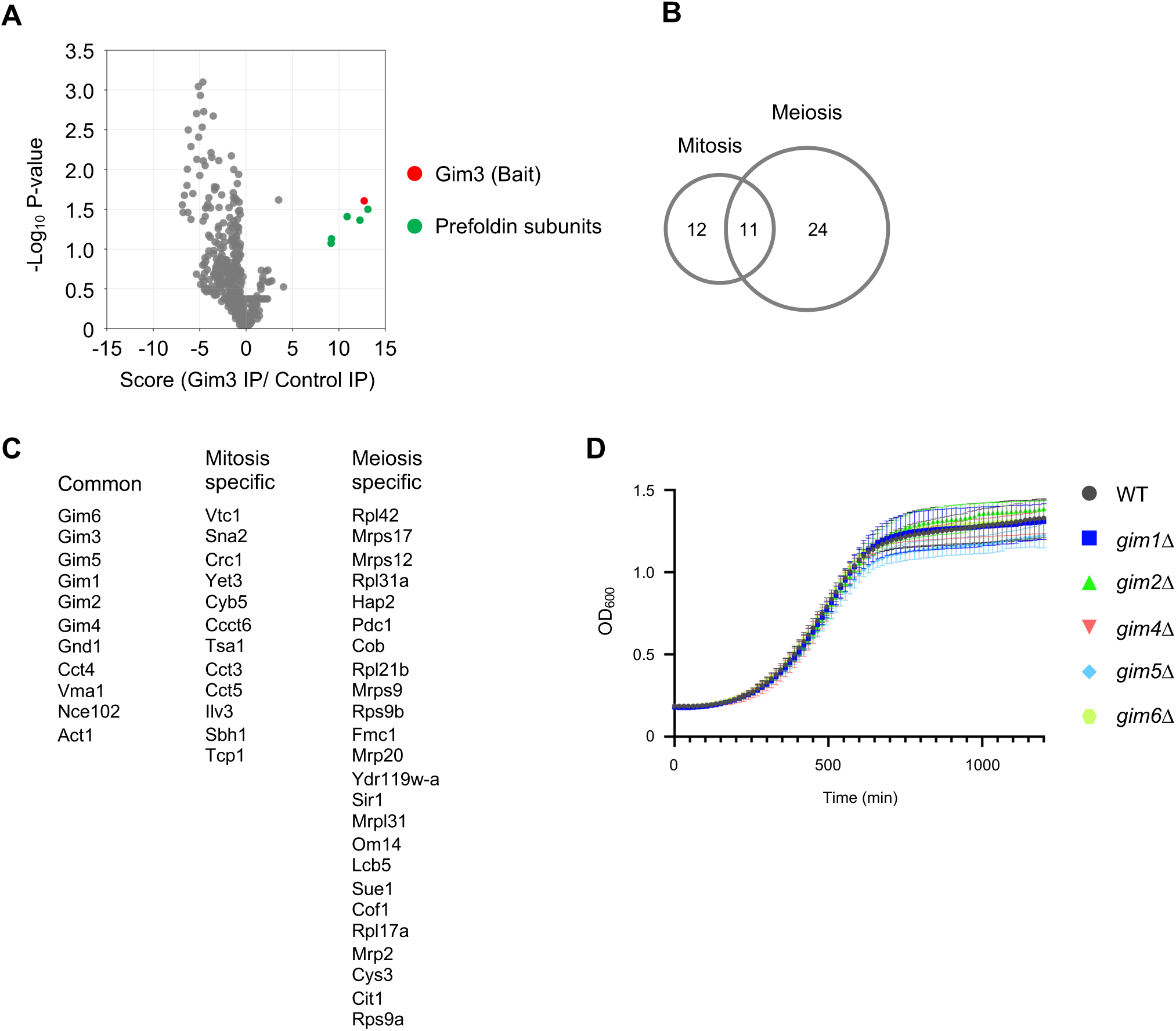
Identification of Gim3 interactors and functional assessment of prefoldin subunits. (A) Mass spectrometry analysis of immunopurified proteins using Gim3 as bait in mitotic cells. The plot shows enrichment scores (x-axis, Gim3 IP versus control IP) versus –log₁₀ P-value (y-axis). Gim3 itself is highlighted in red, and other prefoldin subunits are highlighted in green. (B) Venn diagram showing the overlap of proteins identified in mitotic and meiotic (5 hours in SPO). (C) List of proteins identified in the co-purification experiments under mitotic and meiotic conditions. Proteins are grouped as common, mitosis-specific, or meiosis-specific interactors, corresponding to the Venn diagram in panel B. (D) Growth curves as assessed by OD_600_ measurement of mitotic cells carrying deletions in prefoldin subunits (*gim1Δ*, *gim2Δ*, *gim4Δ, gim5Δ*, and *gim6Δ*) versus WT controls. Growth was monitored in YPD; N = 2-3; data are represented as mean ± SD.

**Supplementary Figure 4.**
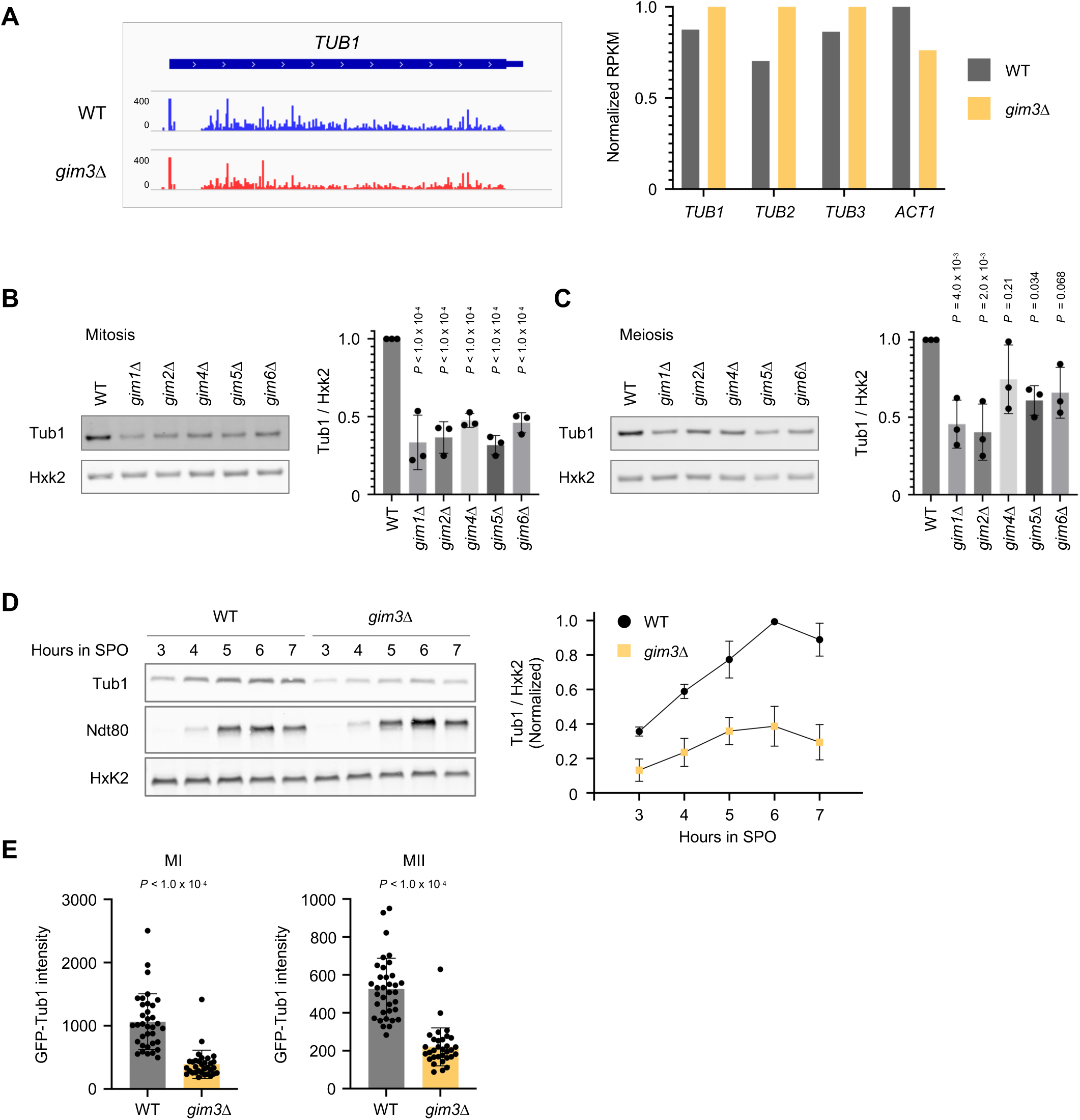
Tub1 protein levels and spindle dynamics in prefoldin subunit deletion mutants. (A) Ribosome profiling analysis during meiosis (4 hours in SPO). Positional ribosome footprint profiles are shown for *TUB1*, and normalized read counts are shown for *TUB1*, *TUB2*, *TUB3*, and *ACT1* in WT and gim3Δ cells. (B) Western blot analysis of Tub1 protein levels in mitotic cells of WT and individual prefoldin subunit deletion strains (*gim1Δ*, *gim2Δ*, *gim4Δ, gim5Δ*, and *gim6Δ*). Hxk2 was used as a loading control. N = 3; data are represented as mean ± SD; Dunnett’s test. (C) Tub1 protein levels in WT and individual prefoldin subunit deletion strains (*gim1Δ*, *gim2Δ*, *gim4Δ*, *gim5Δ*, and *gim6Δ*) strains after 4 hours in SPO (meiosis). Hxk2 was used as a loading control. N = 3; data are represented as mean ± SD; Dunnett’s test. (D) Representative western blot of Tub1 in WT and *gim3Δ* cells during meiosis (left). Hxk2 (Hexokinase isoenzyme 2) was used as a loading control and Ndt80 was used as a marker of meiotic progression. Quantification of Tub1 protein levels (right). N = 3; data are represented as mean ± SD. (E) Quantification of GFP-Tub1 intensity in WT and *gim3Δ* cells during MI and MII. Each dot represents a single cell. Bars indicate mean ± SD. Mann-Whitney U test; 32-35 cells.

**Supplementary Figure 5.**
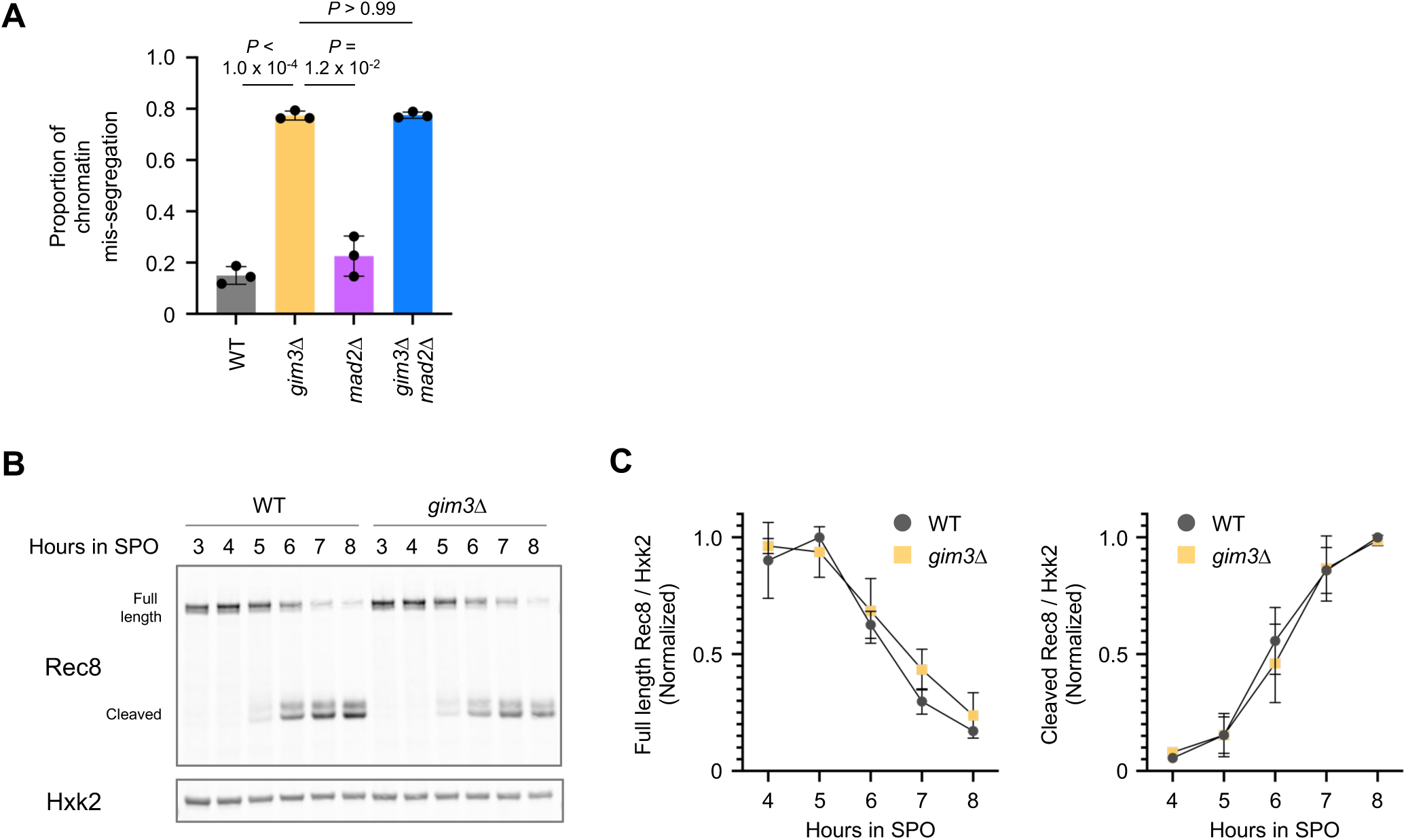
Rec8 cleavage dynamics and checkpoint dependence of mis-segregation in *gim3Δ* cells. (A) Quantification of chromatin mis-segregation in WT, *gim3Δ*, *mad2Δ*, and *gim3Δ mad2Δ* cells. N = 3; data are represented as mean ± SD; Welch’s t-test. (B) Representative western blot of Rec8 in WT and *gim3Δ* cells undergoing meiosis. Deletion of *UBR1*, which stabilizes the cleaved Rec8 fragment, was used to facilitate detection of the Rec8 cleavage product. Hxk2 was used as a loading control. (C) Quantification of uncleaved (left) and cleaved (right) Rec8 protein levels from (B). N = 3; data are represented as mean ± SD.

**Supplementary Figure 6.**
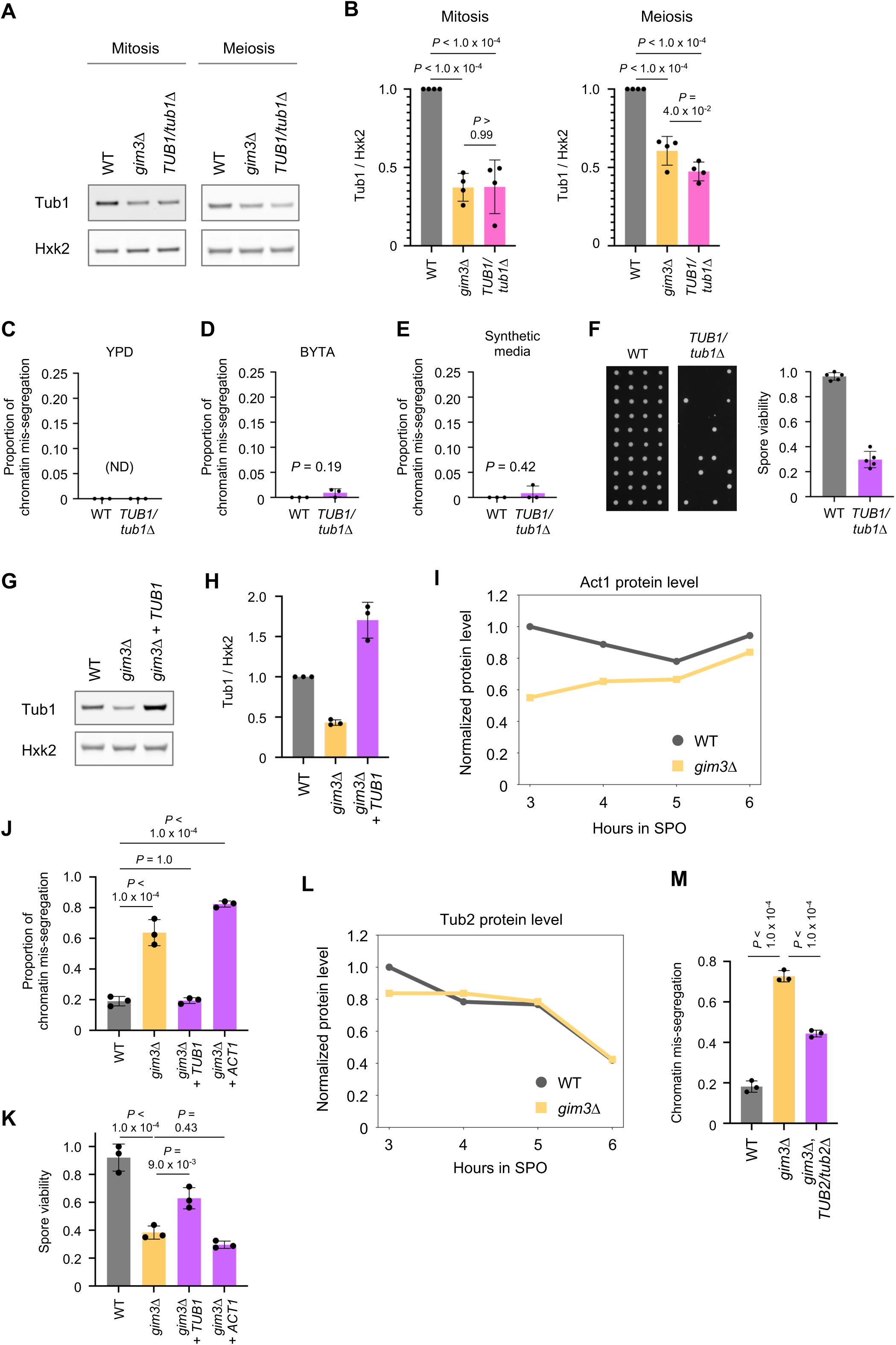
Reduced Tub1 levels or inhibition of tubulin polymerization leads to chromosome mis-segregation. (A) Representative western blot of Tub1 protein levels in WT, *gim3Δ*, and *TUB1/tub1Δ* strains during mitotic growth (left) and meiosis (4 hours in SPO) (right). Hxk2 was used as a loading control. (B) Quantification of Tub1 protein levels from (A). Data are represented as mean ± SD; N = 3; Tukey’s multiple comparison test. (C) Quantification of chromatin mis-segregation during mitosis in WT and *TUB1/tub1Δ* cells in YPD. N = 3; ND, not detected; Welch’s t-test; 64-86 cells were quantified per replicate. (D) Quantification of chromatin mis-segregation during mitosis in WT and *TUB1/tub1Δ* cells in BYTA. N = 3; data are presented as mean ± SD; Welch’s t-test; 66-104 cells were quantified per replicate. (E) Quantification of chromatin mis-segregation during mitosis in WT and *TUB1/tub1Δ* cells in synthetic media. N = 3; data are presented as mean ± SD; Welch’s t-test; 24-44 cells were quantified per replicate. (F) Spore colony growth of tetrads derived from WT and *Tub1/tub1Δ* strains on YPD after 2 days at 30 °C (left). Quantification of spore viability (right). N = 3; data are represented as mean ± SD; Welch’s t-test. 80 spores were quantified per replicate. (G) Representative western blot of Tub1 protein levels in WT, *gim3Δ* and *gim3Δ* overexpressing *TUB1* strains during meiosis (4 hours in SPO). Hxk2 was used as a loading control. (H) Quantification of Tub1 protein levels from (G). Data are represented as mean ± SD; N = 3. (I) Normalized Act1 protein levels in WT and *gim3Δ* cells at 3, 4, 5 and 6 hours in SPO, quantified by mass spectrometry. Mass spectrometry analysis was performed once for each time point. (J) Quantification of chromatin mis-segregation events in WT cells, *gim3Δ* cells carrying the empty cassette, and *gim3Δ* cells expressing *TUB1* or *ACT1*, based on live-cell imaging of Htb1-mCherry. *TUB1* or *ACT1* expression was driven by a GAL promoter and induced using the Gal4-ER system upon addition of β-estradiol (Gao and Pinkham, 2000). β-estradiol was added at 2.5 hour in SPO to a final concentration of 1 nM. N = 3; data are represented as mean ± SD; Tukey’s multiple comparison test. 112-158 cells were quantified per replicate. (K) Quantification of spore viability in WT cells, *gim3Δ* cells carrying the empty cassette, and *gim3Δ* cells expressing *TUB1* or *ACT1*. *TUB1* or *ACT1* expression was induced from a GAL promoter using the Gal4-ER system by addition of 1 nM β-estradiol at 2.5 hour in SPO. Spore viability was determined by tetrad dissection. N = 3; data are represented as mean ± SD; Tukey’s multiple comparison test. (L) Normalized Tub2 protein levels in WT and *gim3Δ* cells at 3, 4, 5 and 6 hours in SPO, quantified by mass spectrometry. Mass spectrometry analysis was performed once for each time point. (M) Quantification of chromatin mis-segregation events in WT, *gim3Δ*, and *gim3Δ TUB2/tub2Δ* cells, based on live-cell imaging of Htb1-mCherry. Chromatin mis-segregation was scored using Htb1-mCherry. N = 3; data are represented as mean ± SD; Tukey’s multiple comparison test. 101-189 cells were quantified per replicate.

**Supplementary Figure 7.**
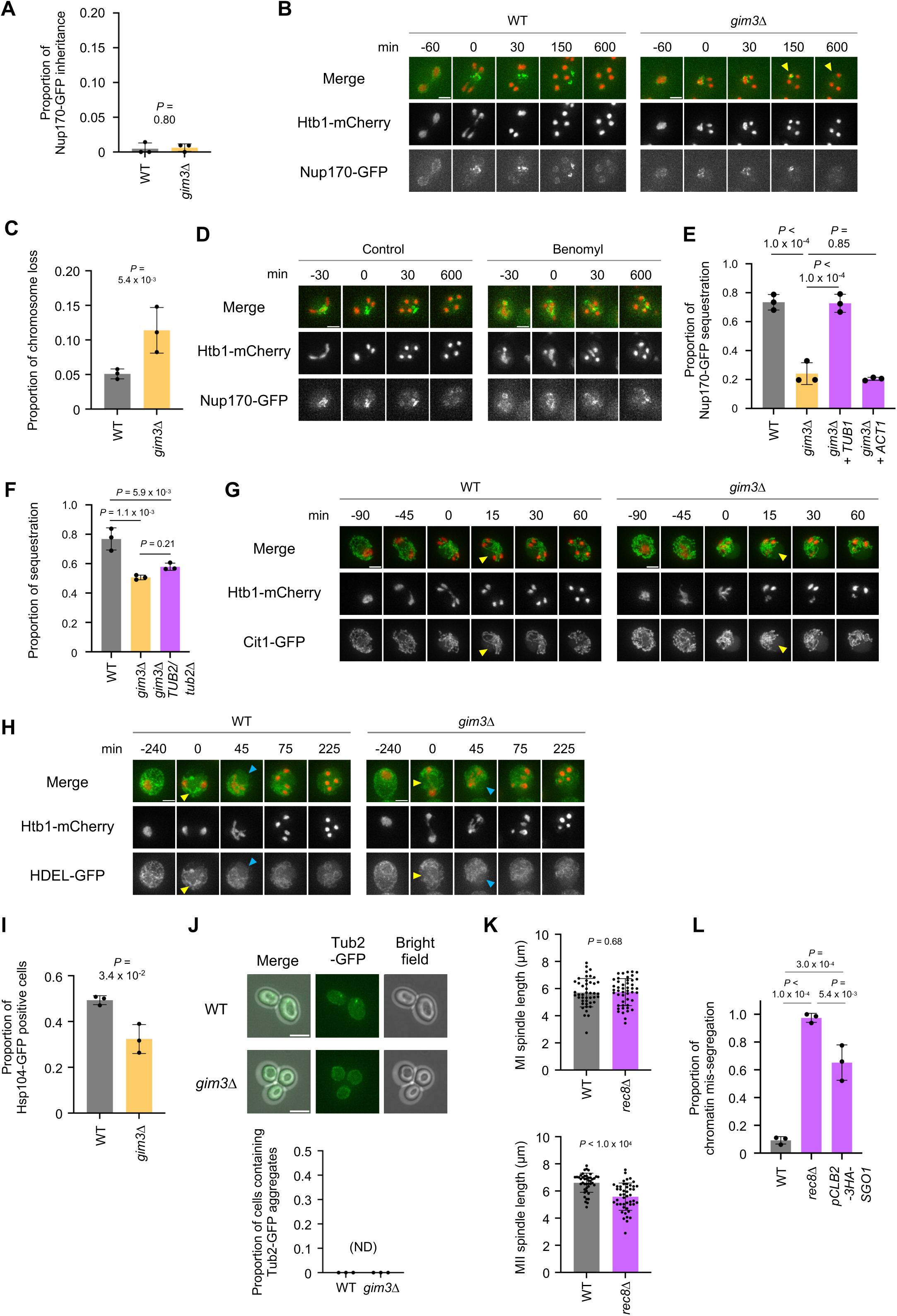
Nuclear component inheritance and organelle remodeling in *gim3Δ* cells during meiosis. (A) Quantification of Nup170-GFP inheritance at 21 hours in SPO. N = 3; data are represented as mean ± SD; t-test. 83-140 cells were quantified per replicate. (B) Live-cell imaging of WT and *gim3Δ* cells expressing Htb1-mCherry and Nup170-GFP during meiosis. Yellow arrowheads indicate degradation of chromatin mass with Nup170-GFP. Time is shown relative to the onset of anaphase II (0 min). Scale bars, 3 µm. (C) Quantification of degradation of chromatin mass with Nup170-GFP shown in (B). N = 3; data are represented as mean ± SD; Welch’s t-test. 63-115 cells were quantified per replicate. (D) Live-cell imaging of WT cells expressing Htb1-mCherry and Nup170-GFP during meiosis treated with benomyl (20 μg/mL). Time is shown relative to the onset of anaphase II (0 min). Scale bars, 3 µm. (E) Quantification of Nup170-GFP sequestration from chromatin in WT cells, gim3Δ cells carrying the empty cassette, and *gim3Δ* cells expressing *TUB1* or *ACT1*. *TUB1* or *ACT1* expression was induced from a GAL promoter using the Gal4-ER system by addition of 1 nM β-estradiol at 2.5 hour in SPO. The WT, *gim3Δ* empty-cassette, and *gim3Δ* cells expressing *TUB1* data are the same as those presented in Figure 5B and are included here for comparison. N = 3; data are represented as mean ± SD; Tukey’s multiple comparison test. 56-88 cells were quantified per replicate. (F) Quantification of Nup170-GFP sequestration from chromatin in *WT*, *gim3Δ*, and *gim3Δ TUB2/tub2Δ* heterozygous diploid cells. N = 3; data are represented as mean ± SD; Tukey’s multiple comparison test. 77-117 cells were quantified per replicate. (G) Live-cell imaging of WT and *gim3Δ* cells expressing Htb1-mCherry and Cit1-GFP (mitochondrial marker) during meiosis. Time is shown relative to the onset of anaphase II (0 min). Yellow arrowheads indicate detachment of mitochondria from the cell periphery. Scale bars, 3 µm. (H) Live-cell imaging of WT and *gim3Δ* cells expressing Htb1-mCherry and HDEL-GFP (ER marker) during meiosis. Time is shown relative to the onset of anaphase II (0 min). Yellow arrowheads indicate the presence (WT) or absence (*gim3Δ*) of ER cables, and blue arrowheads indicate ER detachment from the cell periphery. Scale bars, 3 µm. (I) Quantification of the percentage of Hsp104-GFP positive cells in WT and *gim3Δ* cells at 4 hours in SPO. Hsp104-GFP positive cells were scored based on the presence of visible Hsp104-GFP foci. N = 3; data are represented as mean ± SD; Welch’s t-test. 120-189 cells were quantified per replicate. (J) Representative images of Tub2-GFP in WT and *gim3Δ* cells at 0 hours in SPO (top) and quantification of the percentage of cells containing Tub2-GFP aggregates (bottom). Scale bars, 5 µm. N = 3; data are represented as mean ± SD. ND, not detected. More than 30 cells were quantified per replicate. (K) Quantification of spindle length in WT and *rec8Δ* cells during meiosis I (top) and meiosis II (bottom), measured using Spc42 as a spindle pole body marker. Spindle length was calculated as the distance between Spc42 signals. Data are represented as mean ± SD; Mann-Whitney U test. 44-47 cells were quantified per replicate. (L) Quantification of chromatin mis-segregation events in WT, *rec8Δ*, and *pCLB2-3HA-SGO1* cells, based on live-cell imaging of Htb1-mCherry. Chromatin mis-segregation was scored using Htb1-mCherry as a chromatin marker. N = 3; data are represented as mean ± SD; Tukey’s multiple comparison test. 45-109 cells were quantified per replicate.

**Supplementary Figure 8.**
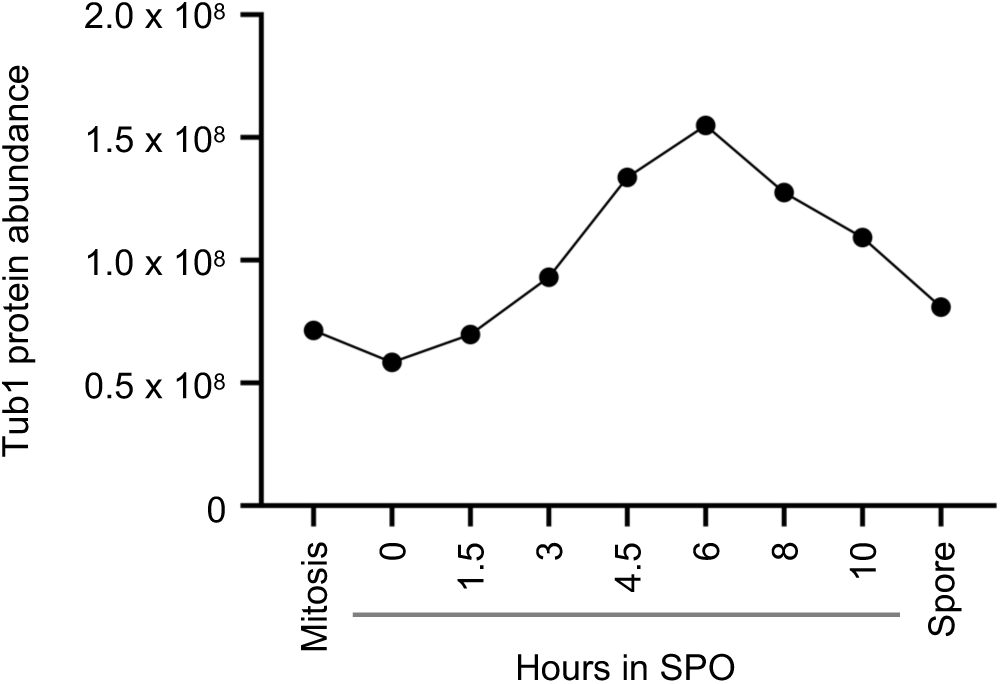
Tub1 protein level during meiosis. Tub1 protein abundance during mitosis and meiosis based on mass spectrometry data reanalyzed from (Cheng et al., 2018).

**Supplementary Figure 9.**
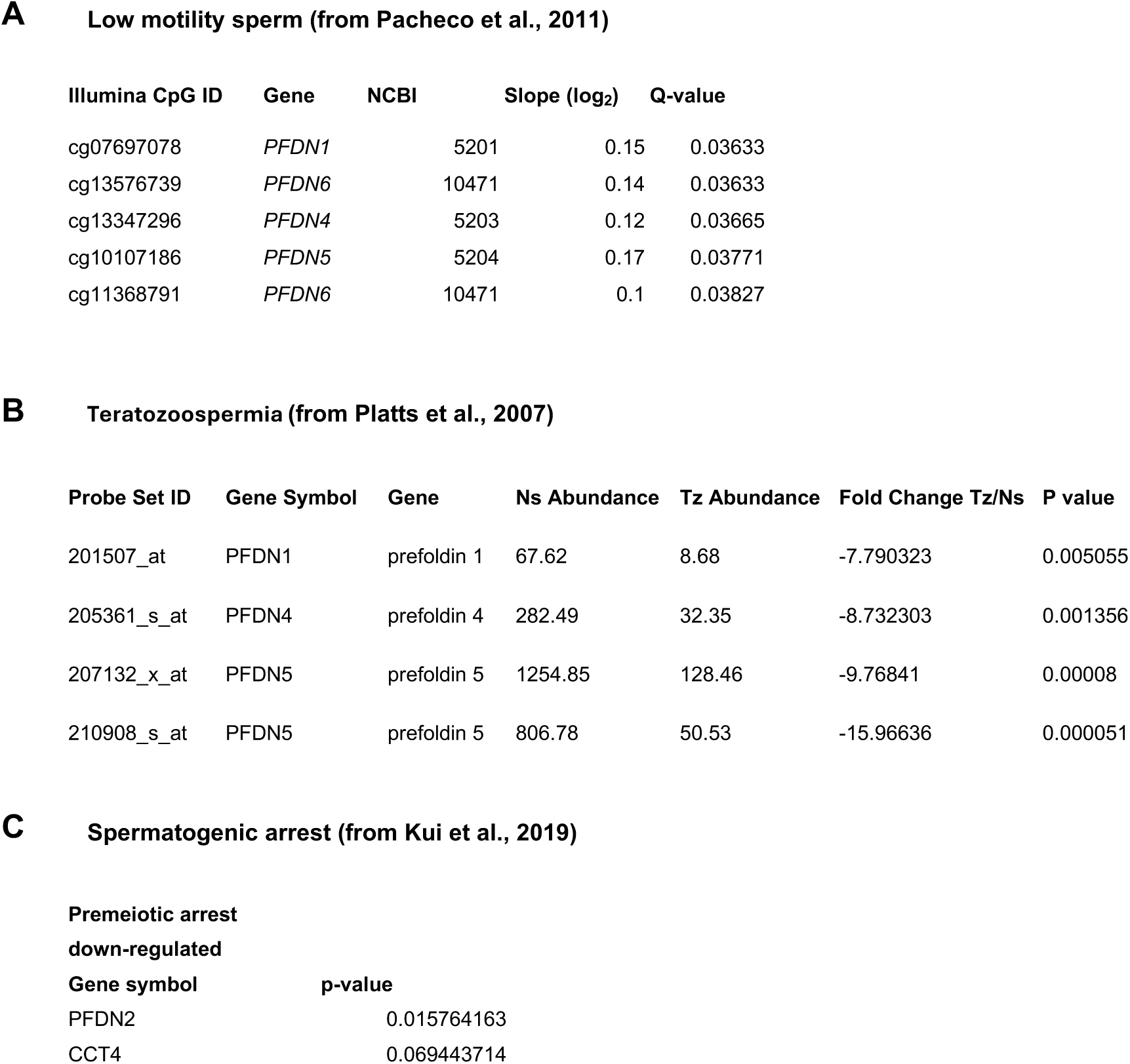
Reanalysis of the association between prefoldin and human phenotypes. Data from (Pacheco et al., 2011) (A), (Platts et al., 2007) (B), and (Kui et al., 2019) (C) were reanalyzed to assess associations between prefoldin and human phenotypes.

**Supplementary Figure 10.**
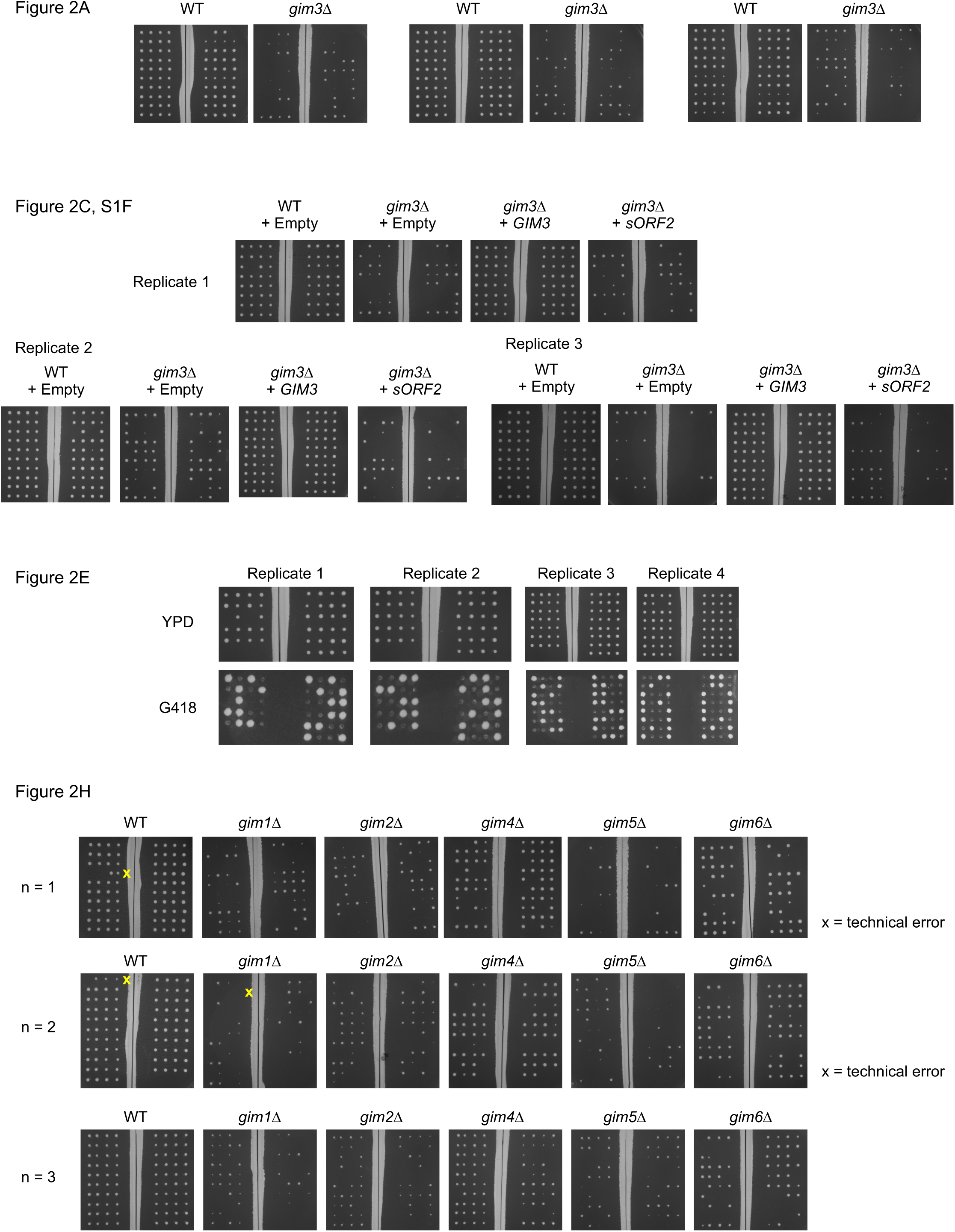
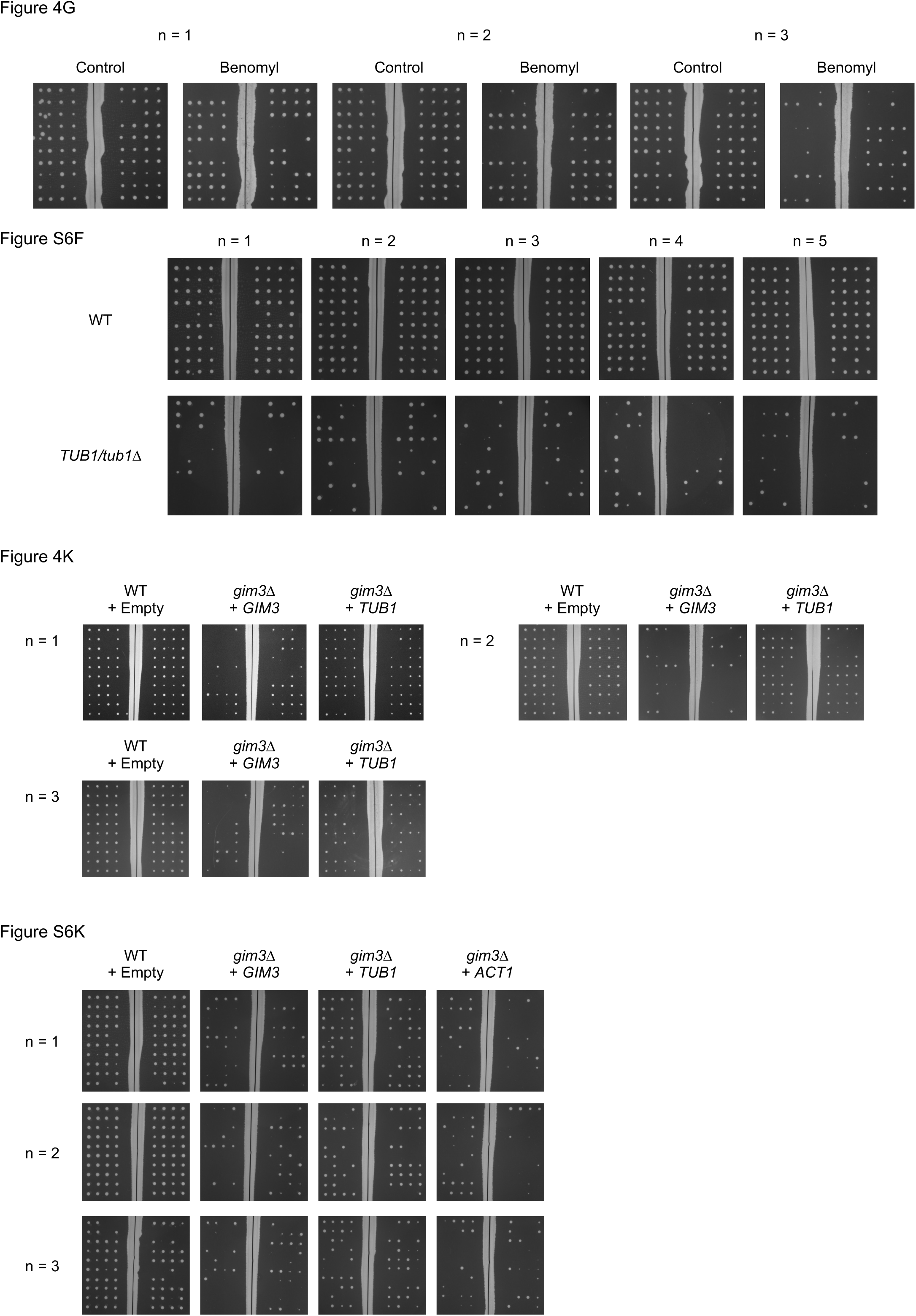
Replicated experiments and full images of gels.

## Supplementary Files

**File S1. CRISPEY-based screening to identify functional sORF loci during meiosis**

This file includes the library sequences (Tab 1), MAGeCK scores (Tab 2), and the positive control genes (Tab 3).

**File S2. Mass spectrometry of Gim3 associated protein levels in mitosis or meiosis**

Cell extracts were immunopurified from Gim3-3V5 cells, and protein abundance was quantified by TMT10-based measurement. Analysis was performed using Spectrum Mill software. Data from three biological replicates for each condition are shown.

**File S3. Mass spectrometry of total protein levels meiotic cells in WT and *gim3Δ***

Cell extracts from WT and *gim3Δ* cells were subjected to mass spectrometry, and protein abundance was quantified by TMT10-based measurement.

**File S4. Ribosome profiling of meiotic cells in WT and *gim3Δ***

RPKM values for WT and *gim3Δ* cells during meiosis (4 hours in SPO).

**File S5. Strains, plasmids, and primers**

This file includes all strains used in this study (Tab 1), as well as plasmids (Tab 2), and primers (Tab 3).

## Full gel images

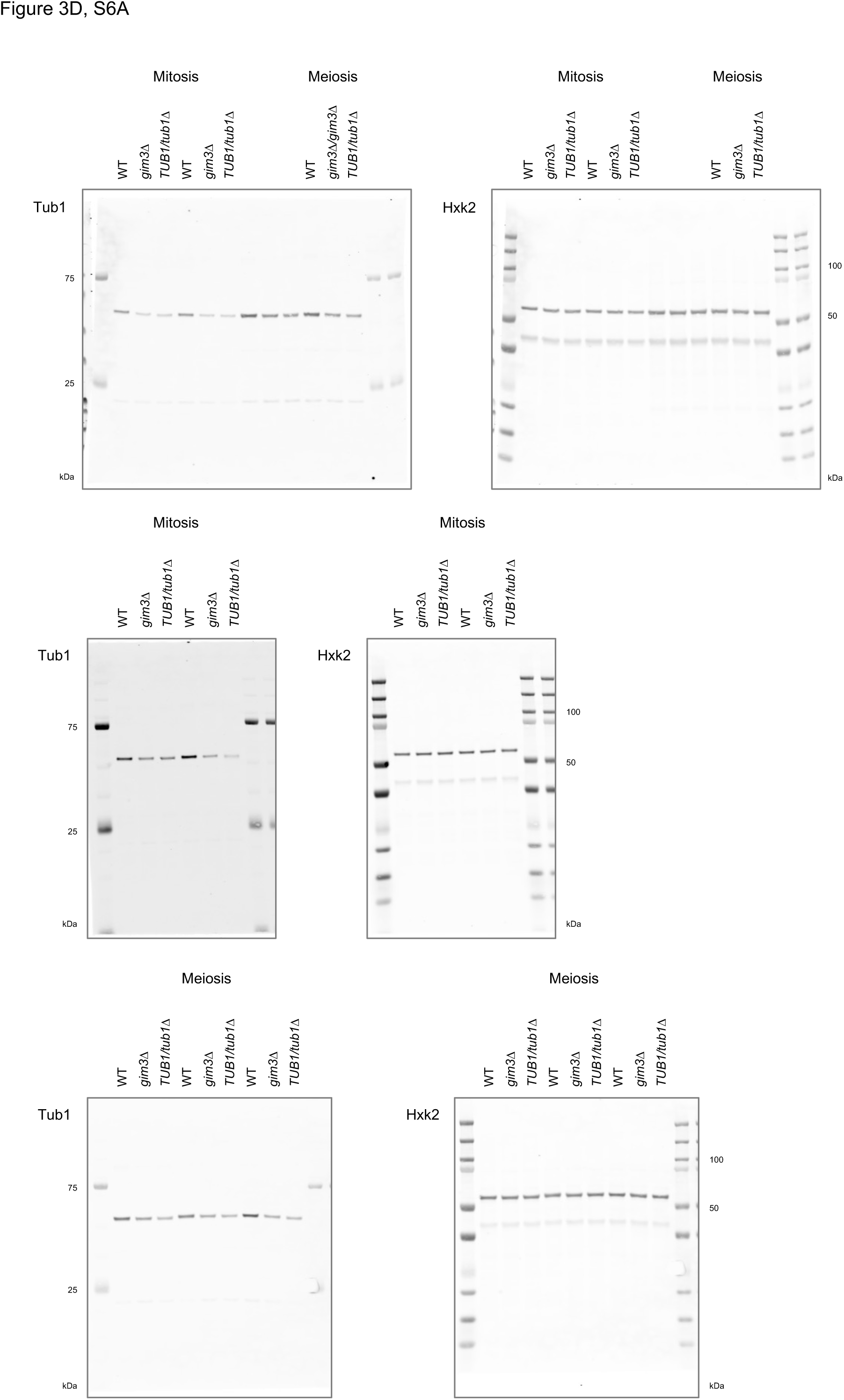

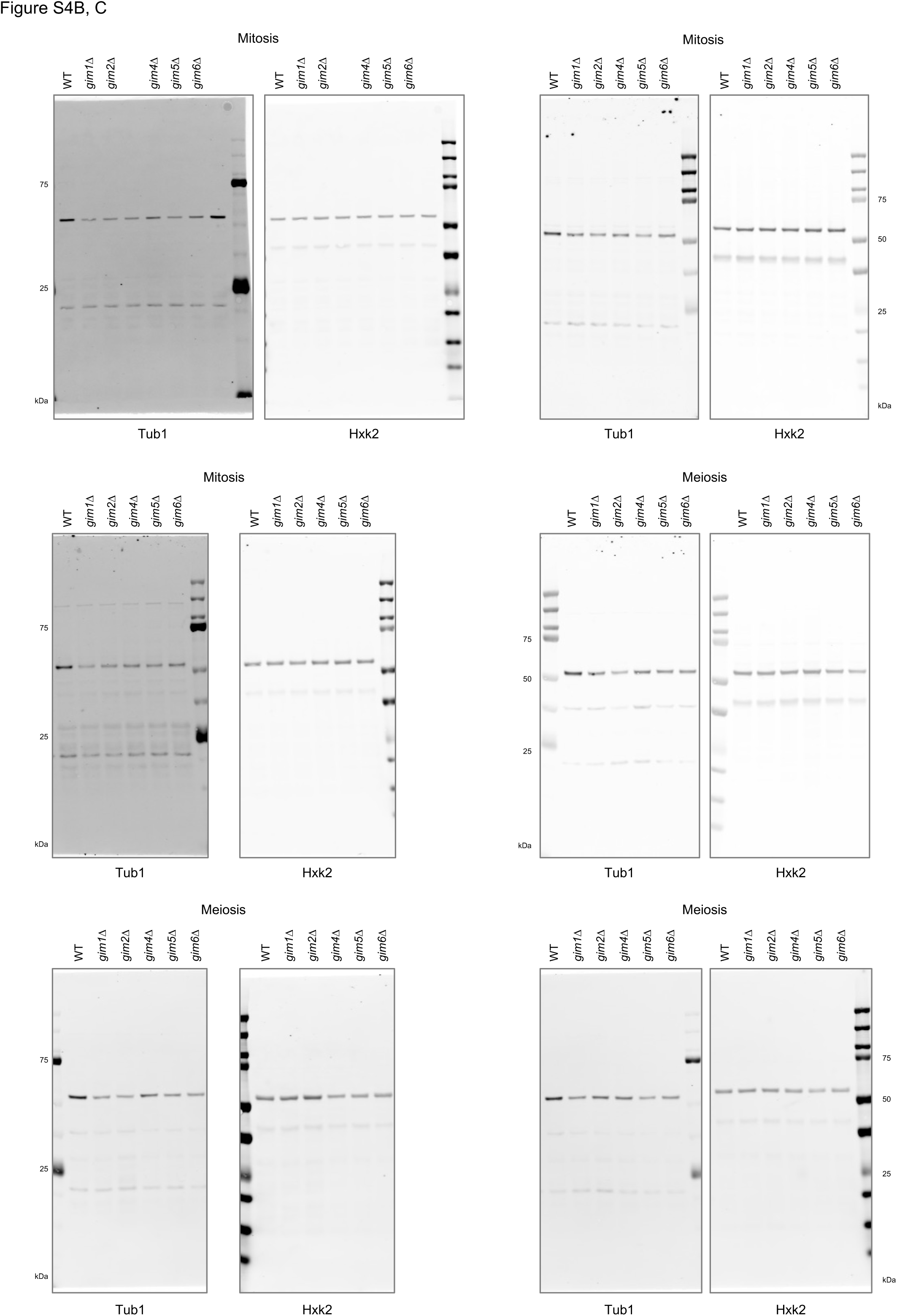

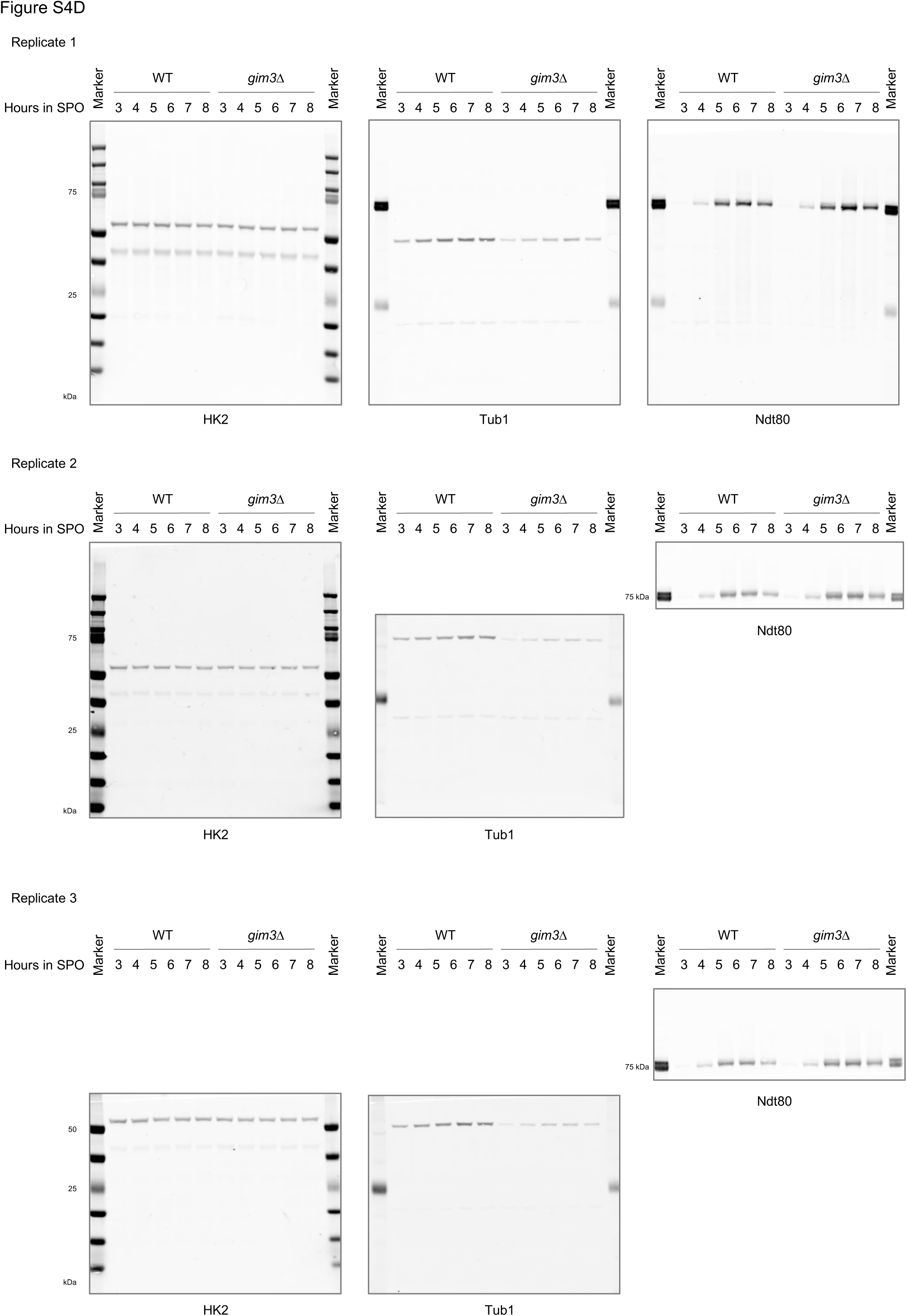

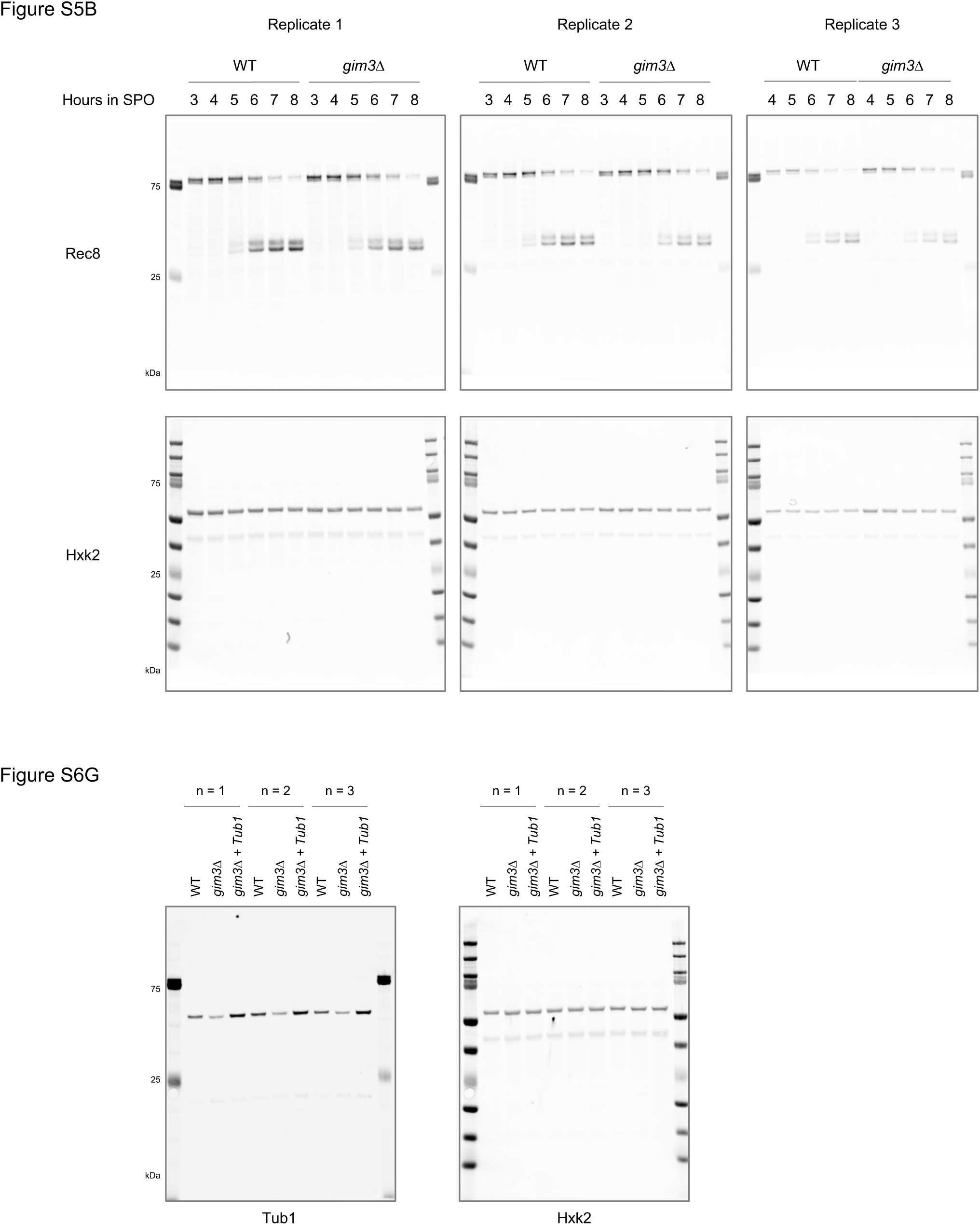

## Notes

### Competing Interest Statement

The authors have declared no competing interest.

